# Optimisation and Pre-clinical Demonstration of Temporal Diffusion Ratio for Imaging Restricted Diffusion

**DOI:** 10.1101/2022.07.25.500826

**Authors:** William Warner, Marco Palombo, Renata Cruz, Noam Shemesh, Derek K. Jones, Flavio Dell’Acqua, Andrada Ianus, Ivana Drobnjak

**Author notes:** shared first author (William Warner, Marco Palombo). shared senior author (Flavio Dell’Acqua, Andrada Ianus, Ivana Drobnjak).

## Abstract

Temporal Diffusion Ratio (TDR) is a recently proposed dMRI technique (Dell’Acqua, 2019) which provides contrast between areas with restricted diffusion and areas either without restricted diffusion or with length scales too small for characterisation. Hence, it has a potential for mapping pore sizes, in particular large axon diameters or other cellular structures. TDR employs the signal from two dMRI acquisitions obtained with the same, large, b-value but with different diffusion times and gradient settings. TDR is advantageous as it employs standard acquisition sequences, does not make any assumptions on the underlying tissue structure and does not require any model fitting, avoiding issues related to model degeneracy. This work for the first time optimises the TDR diffusion sequences in simulation for a range of different tissues and scanner constraints. We extend the original work (which considers substrates containing cylinders) by additionally considering the TDR signal obtained from spherical structures, representing cell soma in tissue. Our results show that contrasting an acquisition with short gradient duration and short diffusion time with an acquisition with long gradient duration and long diffusion time improves the TDR contrast for a wide range of pore configurations. Additionally, in the presence of Rician noise, computing TDR from a subset (50% or fewer) of the acquired diffusion gradients rather than the entire shell as proposed originally further improves the contrast. In the last part of the work the results are demonstrated experimentally on rat spinal cord. In line with simulations, the experimental data shows that optimised TDR improves the contrast compared to non-optimised TDR. Furthermore, we find a strong correlation between TDR and histology measurements of axon diameter. In conclusion, we find that TDR has great potential and is a very promising alternative (or potentially complement) to model-based approaches for mapping pore sizes and restricted diffusion in general.

**Highlights:**

- Temporal Diffusion Ratio (TDR) 2-seq approach maps areas with restricted diffusion
- Optimised gradient waveform pair is: long δ + low G and short δ + high G
- If data is noisy calculating TDR using HARDI acquisition subsets increases accuracy
- We demonstrate TDR for the first time pre-clinically in rat spinal cord
- Pre-clinical TDR values are strongly correlated with axon diameter

## 1. Introduction

Characterising neural tissue structure at the microscopic scale can provide important information regarding development, plasticity, ageing, as well as the impact of various diseases that affect the central and/or peripheral nervous systems^1–5^. For example, axon diameter, along with myelin content, is an important property that influences the speed and efficiency of neural communication^6^. Mapping axon diameter can therefore provide insight into basic brain operation, with larger axons being linked to a faster nerve conduction velocity^6,7^, as well as insight into the progression of neuronal diseases that alter axon diameter distribution, such as amyotrophic lateral sclerosis^8,9^. On the other hand, cell body (namely soma) sizes are of clinical interest in a range of different conditions: for instance, a decrease in neuronal soma size has been reported in subjects with bipolar disorder^10^, an increase in motor neuron soma size is present in amyotrophic lateral sclerosis^11^, while large balloon cells are present in focal cortical dysplasia^12^.

Because of these relationships between tissue microstructure and function in the nervous system, techniques for mapping restricted diffusion in the brain and the corresponding characteristic length-scales are of high potential clinical significance. Non-invasive techniques for measuring brain microstructure, such as those developed using diffusion-weighted MRI (dMRI) are especially of interest, as they provide clinically relevant information whilst obviating the need for invasive biopsy and associated risk.

dMRI sensitises the signal to the displacement of the water molecules in the tissue and provides indirect information about the microscopic tissue organisation^13–15^. Since dMRI techniques provide only an indirect assessment of the tissue microstructure, they often require modelling strategies to inform in a quantitative way on the underlying microstructure. Towards this goal, several methods have been developed in the past to characterise restricted diffusion in tissue and to map cellular sizes, both in white matter (WM) and gray matter (GM).

In white matter, mapping axon diameter has been the focus of several dMRI studies over the last decades^16–32^. Some approaches employ biophysical models of the tissue, for example, representing intra-axonal signal as diffusion restricted in infinitely-long non-permeable cylinders, and extra-axonal signal as Gaussian hindered diffusion^17,18^. Then, the tissue model is fitted to a rich dMRI acquisition, usually with several diffusion-weighting amplitudes (‘shells’) and diffusion times for each shell, to estimate summary statistics drawn from the apparent axon diameter distributions. Another recently proposed approach is to use diffusion measurements with ultra-high diffusion weighting (e.g. b > 6000 s/mm^2^) to map axon diameter sizes based on the departure from the power law expected for diffusion inside cylinders with negligible diameters (i.e. ‘sticks’), however this approach requires at least one b-value much greater than 10 ms/μm^2^, leading to lower resolution data on account of longer TEs and the lower SNR at these high b-values^20^. To improve sensitivity to axon diameter^33^, other approaches go beyond the conventional single diffusion encoding acquisitions and also use oscillating gradient waveforms that can access much shorter diffusion times, either included in a modelling framework^21,34^ or used to contrast measurements with different diffusion times and frequencies^19,35,36^. It has been shown in multiple works^33,37,38^ that measurements of axon diameters and/or diameter distributions are extremely challenging, regardless of the dMRI sequence used, especially when considering standard gradient amplitudes available on clinical scanners. The biggest challenge is the low inherent sensitivity of the signal towards axons of small diameter, with the vast majority of contrast being produced by large axons from the tails of most biological axonal distributions.

In gray matter, mapping cell soma sizes is a more recent endeavour in dMRI, with several studies showing the benefits of including cell soma in microstructure models^39–42^, suggesting mapping restricted spaces such as cell soma is a good additional target for new microstructure imaging techniques. However, it has also been recently shown that, compared to WM, GM may be characterised by faster multi-compartmental exchange of water molecules^43–45^, which represents an extra challenge for the biophysical modelling of dMRI measurements and the unbiased estimation of restricted diffusion in GM.

Furthermore, studying the time dependence of the diffusion signal, as well as of estimated parameters such as diffusion and/or kurtosis tensors, can provide additional information about the tissue characteristics and inform about the type of diffusion processes. For instance, measuring the diffusion time dependence of the diffusion and kurtosis tensors can help to differentiate between the effects of restricted diffusion and inter-compartmental exchange^43–52^, with various applications for brain and body imaging. Although of great interest, characterising both the time dependence and b-value dependence (e.g. to estimate diffusion and kurtosis tensors) requires many measurements and cannot currently be performed in a practical amount of time for clinical applications.

A common feature of the vast majority of dMRI techniques developed for characterising both WM and GM microstructure is the use of simplistic biophysical models to describe the relationship between the measured dMRI signals and the underpinning tissue microstructure. Imaging markers of histologically relevant features, such as axon diameter, are then inferred by fitting such biophysical models. Although very successful for many applications^53,54^, this paradigm has some major limitations: e.g., it relies on strong assumptions on the relevant features characterising the underlying tissue structure and it is prone to ambiguities of results interpretation due to inherent models’ degeneracies.

To bypass some of these limitations, Dell’Acqua et al.^55^ recently introduced the Temporal Diffusion Ratio (TDR), a model-free dMRI approach to characterise restricted diffusion inside large axons using two b-value shells with different gradient timings. Specifically, TDR employs dMRI measurements with a very high b-value (above 6000 s/mm^2^) to suppress fast and assumingly extra-axonal diffusion, and contrasts the signal from two shells at the same b-value obtained with different diffusion times and gradient settings. The advantages of TDR are that it employs standard PGSE diffusion sequences widely accessible on commercially-available MRI systems without any specialist programming, that it does not make any assumptions on the underlying tissue structure, and that it does not require any model fitting, avoiding issues related to model degeneracy^38,56^. Although there has been only one study done so far using TDR^55^, its results have been extremely promising showing the potential for mapping the spatial distribution of large axons in the living human brain.

In this paper we explore the unique potential of the TDR contrast for mapping restriction in tissue. For the first time we optimise the TDR methodology and demonstrate it in a pre-clinical study. We extend the TDR approach from characterising cylindrical restrictions (e.g. axons, as it was originally introduced in previous work^55^), to include both cylindrical and spherical restrictions (e.g., neurites and cellular bodies), allowing us to understand the TDR values observed in a wide range of tissue types. We first employ simulations to optimise the gradient shapes for mapping a wide range of axon diameter distributions, modelled as cylinders, as well as cell body sizes, modelled as spheres. We also investigate the effect of the Rician noise floor on the TDR metrics and present a strategy for maximising TDR contrast by using a subset of the full gradient directions. Finally, we demonstrate the optimised TDR strategies as well as the relationship between TDR and axon diameter using ex-vivo rat spinal cord data.

## 2. Methods

### 2.1 TDR approach

TDR is computed by contrasting two spherically-averaged dMRI signals with the same b-value, each collected with a different set of diffusion gradient parameters (e.g. diffusion time, gradient strength), following the expression:

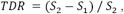

where the sequence parameters used to generate S_1_ and S_2_ are chosen so that S_2_> S_1_ in restricted diffusion. Note that, given that the sequences have the same b-value, Gaussian diffusion would result in equal signal values and no TDR contrast, while in restricted diffusion (and exchange) the specific values of S_1_ and S_2_, as well as TDR contrast, depend on the pore size and diffusion time. In the original implementation of TDR, the two shells are acquired using Single Diffusion Encoding (SDE) sequences, with fixed and equal gradient pulse duration δ, and different diffusion times Δ, one short and one long^55^. Then, the normalised diffusion signals are averaged over gradient directions uniformly sampled over a sphere (e.g. using a HARDI acquisition), and the TDR contrast is calculated based on the powder averaged data, to diminish the effect of fibre orientation distribution:^57^

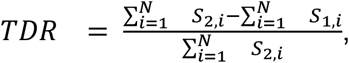

where *N* is the total number of uniformly distributed gradient directions in the HARDI acquisition, *S*_1_,_*i*_ is the signal acquired with the *i*-th gradient direction and gradient waveform used to create *S*_1_, while *S*_2_,_*i*_ is the signal acquired with the *i*-th gradient direction and gradient waveform used to create *S*_2_.

To illustrate the rationale behind the TDR contrast, originally proposed for characterising white matter tissue, we consider a simple tissue model consisting of two compartments: intra-axonal signal is modelled as restricted diffusion inside cylinders and contributes to TDR contrast, and extra-axonal signal is modelled as Gaussian diffusion and does not contribute to the TDR contrast.

Figure 1 shows the signal attenuation and TDR values for spins restricted inside a cylinder as a function of diameter when the diffusion gradients are perpendicular to the fibres. The signals S_1_ and S_2_correspond to sequences with the same b-value (large enough to attenuate extra-cylindrical diffusion) but different gradient waveforms, each yielding a different attenuation for restricted diffusion. For the range of sizes considered here, the difference between S_2_and S_1_, and thus the TDR contrast (right) increases with the cylinder diameter, here proxy for the axon diameter.

**Figure 1.**
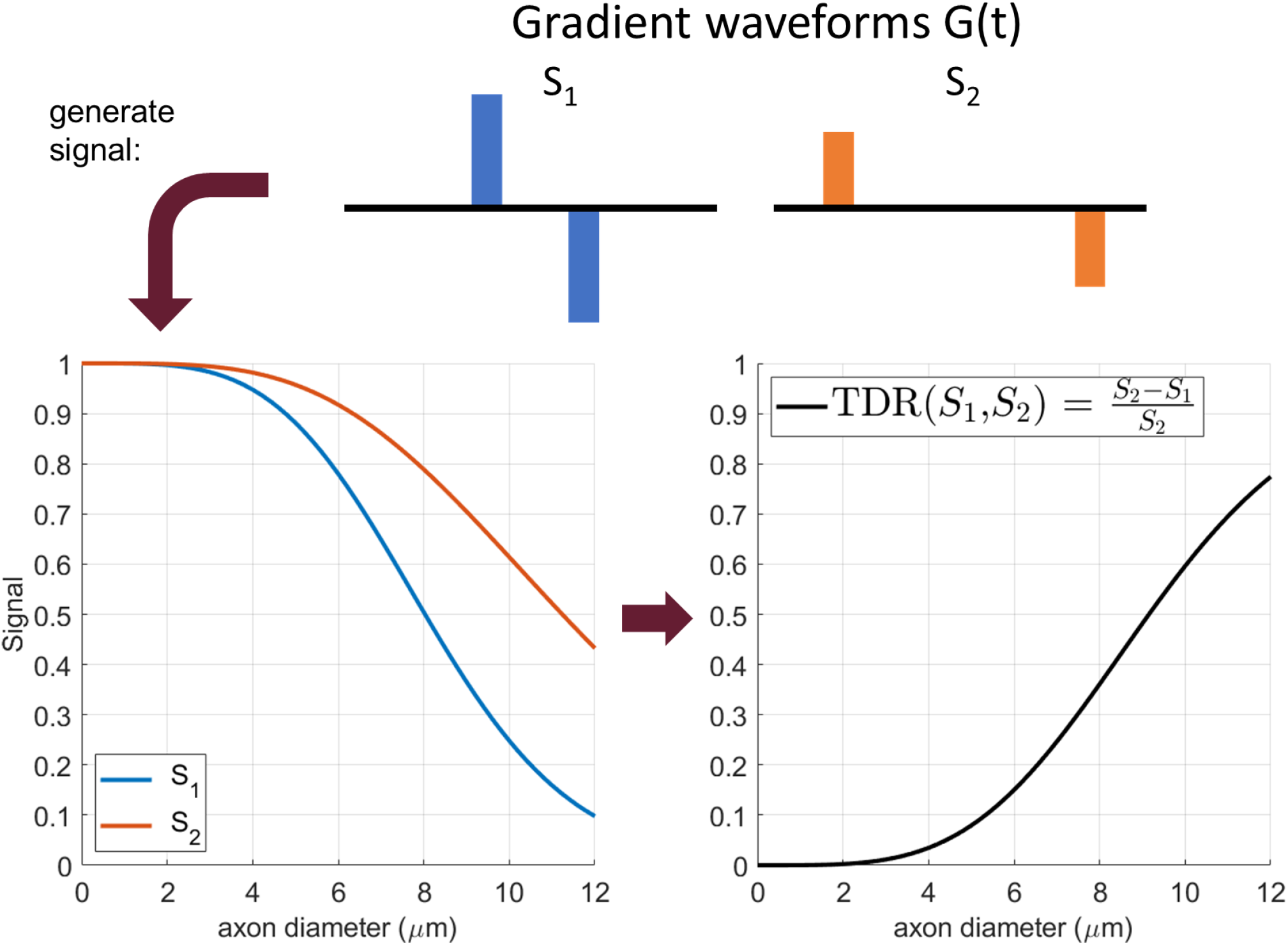
An illustration of the TDR contrast, as proposed in the original study^55^. Both signals (left) are simulated at a b value of 8000 s/mm^2^, with Δ = 21 ms, δ = 9 ms, G = 276.8 mT/m for S_1_ and Δ = 55 ms, δ = 9 ms, G = 162.9 mT/m for S_2_, respectively. The relative difference between the two signals produces the TDR contrast on the right, that is monotonically increasing with pore size for the range considered here.

In reality, each white matter voxel contains a range of axon diameters that contribute to the signals S_1_ and S_2_. In this case the TDR contrast is effectively the integral over the distribution of axon diameters. Hence, given that the difference between S_1_ and S_2_increases with axon diameter, TDR contrast will be higher in voxels containing many large axons. In other words, the TDR contrast is mostly sensitive to large axons and therefore has the potential to map regions where there are larger axons.

In this work, we also analyse the TDR contrast in spherical pores, that have been included as a signal compartment when modelling gray matter^39^ as well as other tissue types, for instance tumours^58,59^.

### 2.2 Simulations

In this study we perform simulations to design optimised SDE acquisitions that maximise TDR contrast. In the first set of simulations we optimise the gradient waveform, then we explore the intensity and the potential of TDR as an imaging contrast.

Simulations are generated using the Microstructure Imaging Sequence Simulation Toolbox (MISST, version 0.93)^60^, a diffusion MRI simulator that calculates the signal attenuation for various restricted geometries and different gradient waveforms using the matrix formalism^61–63^. Code for MISST is available at http://mig.cs.ucl.ac.uk/index.php?n=Tutorial.MISST. Code written by the authors will be made available before the final submission.

#### Simulation 1: Optimising gradient waveforms for TDR

In our first simulation, we optimise the gradient waveform for each of the two SDE sequences to maximise TDR contrast. We consider substrate configurations for two different geometries: a) distributions of parallel impermeable cylinders (mimicking axons), and b) distributions of impermeable spheres (mimicking cell bodies), illustrated in Figure 2. For each of these two geometries, we consider two diameter distributions, one large and one small. For cylinders we use large and small gamma distributions to mimic axons typically found in the spinal cord^24,64^ (mean = 5.33 μm, std = 3.00 μm) and corpus callosum^24,65^ (mean = 1.93 μm, std = 0.81 μm), respectively. For spheres we use large^66,67^ (mean = 15 μm, std = 0.5 μm) and small^67–69^ (mean = 7 μm, std = 0.5 μm) normal distributions to mimic cell somas typically found in mammalian brains. The intrinsic diffusivity for all pores is set to 2 μm^2^/ms. The signal from extracellular space, as well as any exchange effects are assumed to be negligible for the TDR contrast.

**Figure 2.**
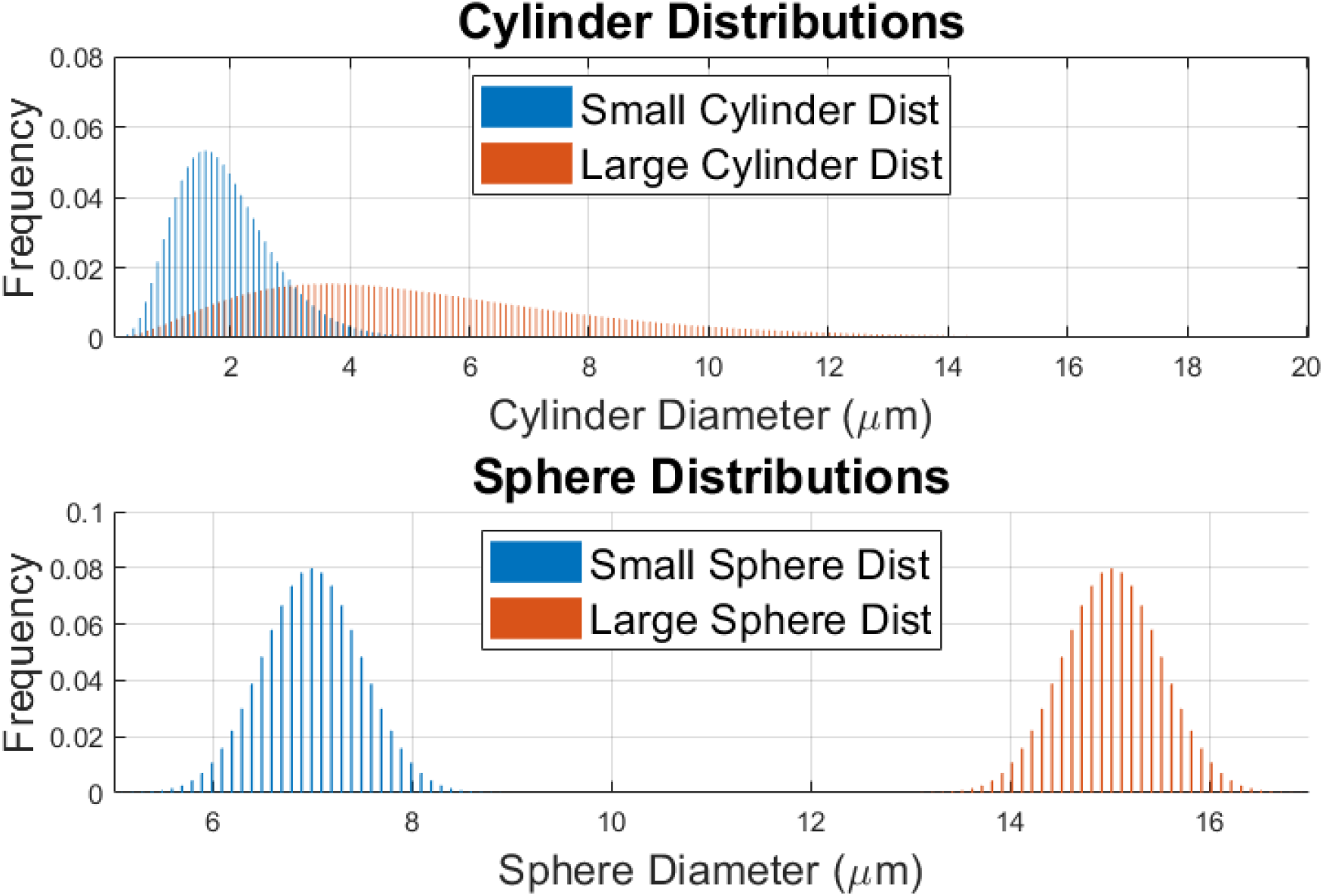
An illustration of the substrate distributions used: two gamma distributions of parallel cylinders (small: mean = 1.93 μm, std = 0.81 μm; large: mean = 5.33 μm, std = 3.00 μm) and two normal distributions of spheres (small: mean = 7 μm, std = 0.5 μm; large: mean = 15 μm, std = 0.5 μm).

We assume the same b-value for both SDE sequences in order to ensure the diffusion contrast from Gaussian diffusion is the same. We also set b to a very high value, specifically b=8000 s/mm^2^ as in the initial work introducing TDR^55^, in order to eliminate the signal that comes from the extracellular space and free water. TDR is calculated from normalised diffusion signals averaged over 60 gradient directions uniformly sampled over a sphere as in the original implementation of TDR^55^.

Optimisation is done using a range of simulations performed with MISST and spanning a large space of sequence parameters, for three high performance animal and human scanner hardware constraints: 1) G < 600 mT/m and Δ+δ < 45 ms corresponding to a typical pre-clinical scanner; 2) G < 2700 mT/m, Δ+δ < 45 ms corresponding to a high-gradient pre-clinical system; 3) G < 300 mT/m, Δ+δ < 80 ms corresponding to the (human MRI) Connectom scanner (Siemens Healthineers). Optimisation is automated using the interior point algorithm as implemented in the MATLAB fmincon function, with Δ, δ and G values for each of the two sequences as linked variables under investigation. The interdependence of these variables means we are ultimately optimising over a 4-dimensional space [Δ_1_, δ_1_, Δ_2_, δ_2_] with Δ and δ for each of the two sequences determining the appropriate G-values. Note that to keep consistent with previous TDR work and widely available pre-clinical and clinical settings, we only considered SDE waveforms (i.e. other waveforms such as double diffusion encoding, oscillating gradients or b-tensor encoding were not considered here).

We then evaluate the performance of the optimised gradient waveforms against a set of gradients chosen according to the previously published strategy for TDR^55^, namely both sequences have the same, short gradient duration, and only the diffusion time is varied: S_2_ has a long diffusion time and S_1_ has a short diffusion time. The exact values of δ, Δ and G were chosen according to the scanner constraints (1-3), with the minimum possible δ and Δ for S_1_ and maximum possible Δ for S_2_. As these sequences were not determined based on full scale optimisation, we will refer to them as “non-optimised” waveforms/sequences.

#### Simulation 2: Effect of restriction size on TDR

In the second simulation we use optimised sequences obtained from simulation 1 to evaluate the TDR contrast over a wide range of pore size distributions.

We consider both distributions of cylinders and spheres to model restrictions in both white matter and gray matter, with size distributions commonly found in tissue^24,39,65–69^. Cylinder sizes follow a gamma distribution^18,24^, whilst sphere sizes follow a normal distribution^66^. For cylinders, the gamma distributions are truncated at 20 μm to match realistic values from the tissue; thus, combinations of parameters where this truncation changes the mean by more than 10% are not considered. For all substrates, intrinsic diffusivity is set to 2 μm^2^/ms and extracellular space is neglected. These simulations are run without noise to illustrate the maximum potential of TDR as an imaging contrast.

#### Simulation 3: Effect of gradient directions and noise on TDR

In the third simulation, we investigate the effect of the Rician noise floor on TDR values. To this end, we consider different fibre configularions: one, two and three fibre bundles consisting of parallel cylinders with the same diameter distributions (Gamma distribution with mean = 5.33 μm, std = 3.00 μm) with separate fibre bundles crossing at right angles (in the case of two and three fibres), as well as spherical pores (Gaussian distribution with mean = 7μm, std = 0.5μm), and we simulate the signals using the optimised sequences from simulation 1. Then, for each measurement, we consider Rician noise at SNR levels of 20 and 50 in the non-diffusion weighted images, typical for for clinical and pre-clinical acquisitions^70,71,^ as well as noise free signals, i.e. SNR ∞.

We also look at the effect of the orientation distribution of gradient directions included when calculating TDR. In the previous work^55^ S_1_ and S_2_ are calculated as the signal average over a set of uniformly distributed gradient directions. However, in white matter, due to the fast signal decay in certain directions (e.g. parallel to the fibres), the contribution to the direction-averaged signal of these measurements will carry little to no extra information about the axon diameter and could potentially introduce bias to the TDR estimation, due to the noise floor. Here we investigate this effect and its impact on TDR.

### 2.3 Preclinical experiments

All animal studies were approved by the competent institutional and national authorities, and performed according to European Directive 2010/63.

The preclinical experiments aim to investigate TDR contrasts in ex-vivo rat spinal cord, to validate our optimisation, both in terms of gradient waveforms as well as the number of HARDI directions employed, and the relationship between TDR and axon diameter in different WM ROIs.

The data and code from the preclinical experiments are available at https://github.com/andrada-ianus/TDR_study.git.

#### Data acquisition

One rat spinal cord was extracted via transcardial perfusion with 4% Paraformaldehyde (PFA). After extraction, the spinal cord was immersed in a 4% PFA solution for 24 h, and then washed in a Phosphate-Buffered Saline (PBS) solution for 24 h. Two sections of cervical spinal cord were cut and placed separately in 5 mm NMR tubes filled with Fluorinert (Sigma Aldrich, Lisbon, PT). The samples were imaged on a 16.4 T Bruker Aeon Ascend scanner (Bruker, Karlsruhe, Germany) equipped with a 5 mm birdcage coil and gradients capable of producing up to 3 T/m in all directions.

#### Imaging protocol

Diffusion MRI datasets for TDR were acquired using a SDE-EPI sequence with the following parameters: TE = 49 ms, TR = 2 s, 16 averages, slice thickness = 0.5 mm, 5 slices, in plane resolution = 0.09 × 0.09 mm^2^, matrix = 64 × 64, Partial Fourier = 1.1. The EPI acquisition bandwidth was 400 kHz and data was acquired in a single shot with a total acquisition time of 1h50m.

In terms of diffusion weighting, the TDR acquisition was performed for a fixed b-value of 8000 s/mm^2^, similar to the protocol originally proposed in Dell’Acqua^55^, in two scenarios: 1) the sequence parameters were chosen so that the maximum gradient strength was limited to 600 mT/m, a value available on many preclinical systems, and 2) the maximum gradient strength was 2500 mT/m, close to the maximum available on this gradient system. For each scenario, we considered waveforms with the same gradient duration and different diffusion times, as originally proposed in^55^, referred to as the non-optimised protocol, as well as the optimised waveforms proposed in this study. Each shell was acquired with 10 b0 values and 60 diffusion directions each, and the specific timing parameters for the non-optimised and optimised protocols are provided in section 3.2.

The data was acquired with an in-house implementation of the sequence which loops through the different diffusion times in order to avoid any potential signal differences caused by sequence adjustments. The sequences implemented in PV6.0.1 are available upon request.

#### Data analysis

Pre-processing: Complex data were denoised using the MP-PCA approach^72^ following the steps described in ^71^ to account for the effect of Partial Fourier acquisition in the data, and the magnitude was computed. Then the data was normalised for each shell.

#### ROI analysis

For selected WM tracts with different axonal properties (Vestibulospinal (VST), Reticulospinal (ReST), Rubrospinal (RST), dorsal corticospinal (dCST), Funiculus Cuneatus (FC) and Funiculus Gracilis (FG)), the averaged TDR values were compared with axon diameters estimated from quantitative histology reported in the literature^73^.

## 3. Results

### 3.1 Simulations

#### Simulation 1: Optimising gradient waveforms for TDR

This section presents the optimisation results of the TDR contrast for sequence parameters typical for pre-clinical scanners, namely G < 600 mT/m and Δ+δ < 45 ms. Optimised TDR acquisitions for other scanner settings, namely G < 2700 mT/m, Δ+δ < 45 ms, corresponding to a high-gradient pre-clinical system, and G < 300 mT/m, Δ+δ < 80 ms, corresponding to the Connectom scanner, are shown in Supplementary Information in Figure S1.

Figure 3a illustrates diffusion weighted signal values averaged over 60 uniformly distributed gradient directions for a range of different combinations of gradient durations and diffusion times and a substrate consisting of small cylinders. Signal values ranging from low (blue) to high (red) values are shown, and TDR is calculated for each pair. In order to find the combination of diffusion sequence parameters that maximises TDR, the optimisation looks for values for S_1_ and S_2_ that are most different from one another. The corresponding, optimal, pair of gradient waveform parameters (squares) has one gradient waveform with short duration and diffusion time and the other with long duration and diffusion time. The short duration waveform is the same as the short diffusion time sequence in the “non-optimised” approach (circles), while the second optimal waveform has much longer gradient duration than any sequence in the “non-optimised” approach, where the diffusion duration is kept constant between the two waveforms and only diffusion time has changed.

**Figure 3.**
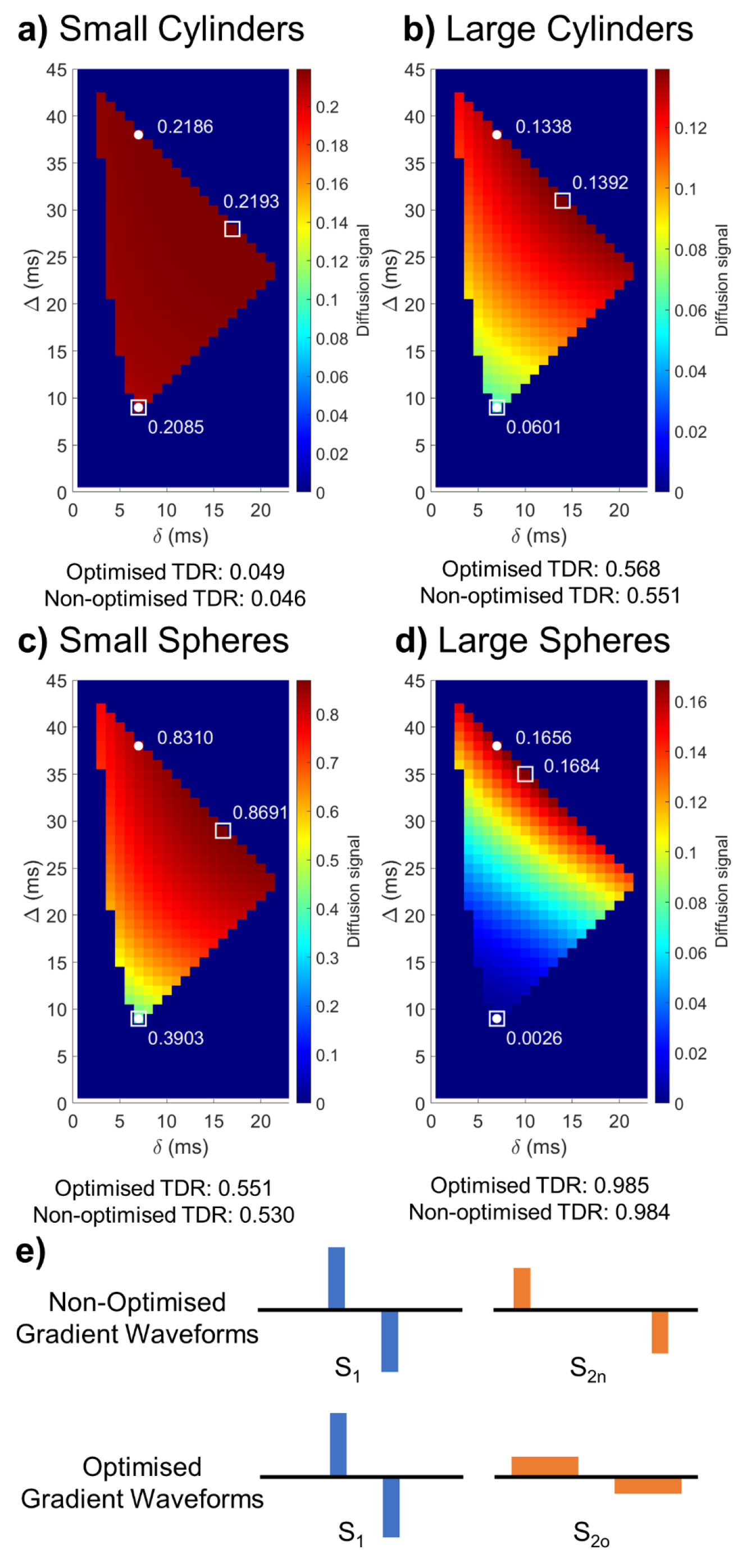
Optimisation results maximising TDR. (a-d) Maps showing the diffusion weighted signal averaged over the 60 uniformly distributed directions for sequences with b=8000 s/mm_2_and various δ/Δ combinations for the substrates illustrated in Figure 2. White markers indicate the non-optimised (circle) and optimised (square) sequences, respectively. The optimised sequences provide larger signal differences between S_1_ and S_2o_compared to between S_1_ and S_2n_, and higher TDR values. (e) Schematic representation of non-optimized and optimised gradient shapes. These figures show that optimised TDR maximises the difference between the signal from the two acquisitions, using sequences with different pulse shapes.

Similar results are obtained for other substrates as well: large cylinders (b), small spheres (c) and large spheres (d) - although for spheres with large diameters the optimal duration of the long gradient sequence is slightly shorter. The exact values for all optimised waveform parameters as determined through non-linear optimisation are:

- small cylinders: S_1_ : Δ=8.87 ms, δ=6.87 ms; S_2o_: Δ=28.46 ms, δ=16.54 ms
- large cylinders: S_1_ : Δ=8.87 ms, δ=6.87 ms; S_2o_: Δ=30.92 ms, δ=14.08 ms
- small spheres: S_1_ : Δ=8.87 ms, δ=6.87 ms; S_2o_: Δ=29.04 ms, δ=15.96 ms
- large spheres: S_1_ : Δ=8.87 ms, δ=6.87 ms; S_2o_: Δ=35.13 ms, δ=9.87 ms

In addition to the specific substrates shown in Figure 3, we have also performed the optimisation for other substrates, both consisting of cylinders and spheres. Whilst distributions that include very large pores can have different optimal sequence shapes, in all cases considered, increasing δ for the S_2o_ shell improves the contrast compared to the non-optimised version of TDR.

#### Simulation 2: Effect of restriction size on TDR

This section explores the TDR contrast over a wide range of pore size distributions, to explore the types of structures visible through TDR. Following the previous results which show that the optimised sequences are very similar across a range of small and large cylinders and spheres, in this simulation we chose the optimised parameters obtained for the large cylinder distribution, thus for S_1_ we choose Δ = 8.87ms, δ = 6.87ms and for S_2_ we choose Δ = 30.92ms, δ = 14.08ms.

Figure 4a illustrates TDR values across substrates consisting of cylinder distribution with a wide range of parameters (mean along the x-axis and standard deviation along the y-axis), calculated based on the optimised sequences above. The plot presents the overall link between the axon distribution sizes and TDR and shows that distributions with the mean below 3μm and standard deviation below 1μm - resolution limit for this gradient strength - have TDR values close to zero, and hence would not be detectable in the images. On the other hand larger axons have sufficiently large TDRs (TDR=1 is a maximum value) which the approach will pick up, and so will stand out in the TDR map images. This plot shows where typical axon distributions found in the tissue would be: “small dist” matches a previous model of callosal white matter^24^ which is undetectable with this approach (as expected based on the max gradient strength used^37^) and “large dist” replicates the white matter structure in the spinal cord, which is within the sensitivity of the TDR approach.

**Figure 4.**
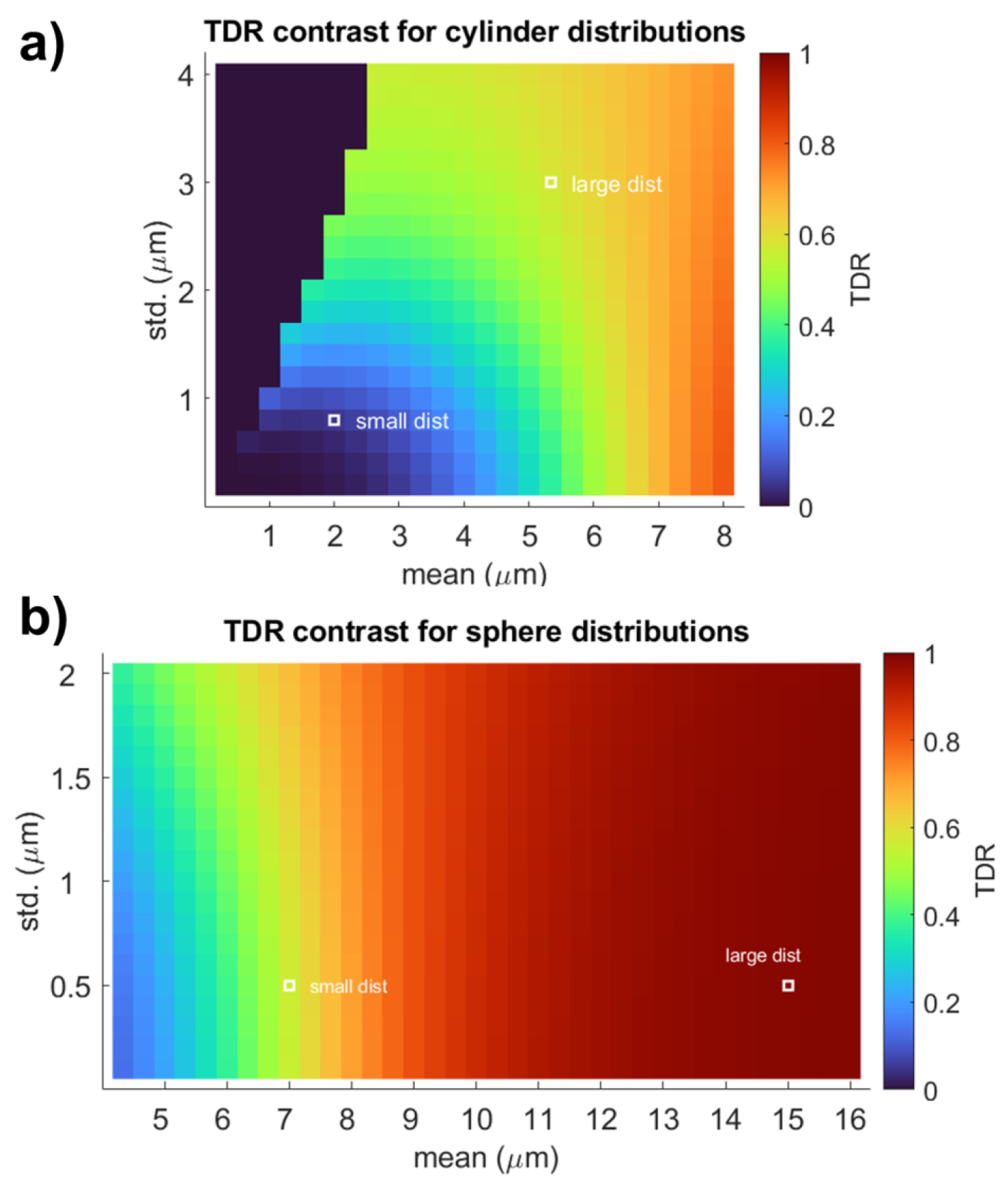
Noise-free TDR values calculated for sequences with optimised parameters for G_max_=600 mT/m: S_1_ : Δ = 8.87 ms, δ = 6.87 ms; S_2o_: Δ = 30.92 ms, δ = 14.08 ms. The signal is simulated across a wide range of (a) cylinder and (b) spherical diameter distributions. For cylinders, we simulate Gamma distributions, and for spheres we simulate Gaussian distributions, which reflect the size distributions usually measured in the tissue. The typical large and small size distributions presented in Figure 2 are shown using white markers. For cylinders, the gamma distributions are truncated at 20 μm to match realistic values from the tissue; thus, combinations of parameters where this truncation changes the mean by more than 10% were not considered.

The isocolors in Figure 4a show existing ambiguities in associating a specific TDR value to a single size distribution. Multiple size distributions can provide the same TDR value: e.g., all the distributions characterised by mean and standard deviation along the green area in Figure 4 would all contribute to a TDR value of ∼0.4. This is expected to be seen in the TDR approach whose main aim is to visualise areas with large, detectable restrictions rather than to map their exact size.

Figure 4b) shows results for spherical substrates. These are much larger as they represent the structures in the gray matter and their TDR values are much higher and, depending on the SNR, would be detectable in the TDR maps for the hardware parameters selected.

#### Simulation 3: Effect of gradient directions and noise on TDR

It is well known that when imaging a single bundle of parallel fibres, the intracellular signal obtained depends on the angle between the orientation of the fibre bundle and the direction of the diffusion sensitising gradients: orthogonal gradients return high signal, whilst parallel gradients create stronger signal attenuation and return lower signal. This is particularly emphasised when the b-values are very high^33^ such as the case in the TDR approach. A similar rationale applies to imaging more than one fibre bundle: the gradient directions returning the highest signals are those which are close to perpendicular to one or more fibre bundles; meanwhile gradient directions which are not close to perpendicular to any of the fibre bundles show very low signal. This concept is represented in Figure 5, which shows signal measurements for two crossing fibre bundles and 60 uniformly distributed gradient directions (Figure 5a). Figure 5b shows all the 60 signal measurements colour coded according to the angle between the plane of the fibres and the gradient direction shown in a) (e.g. red are gradient directions most perpendicular to the fibres). Figure 5c shows all 60 signal measurements ordered from the highest to the lowest signal to emphasise the impact that gradient direction has on the signal itself. It can be seen how the signal measurements are highly distributed from very high, through medium intensity and finally some measurements with very low signal.

**Figure 5.**
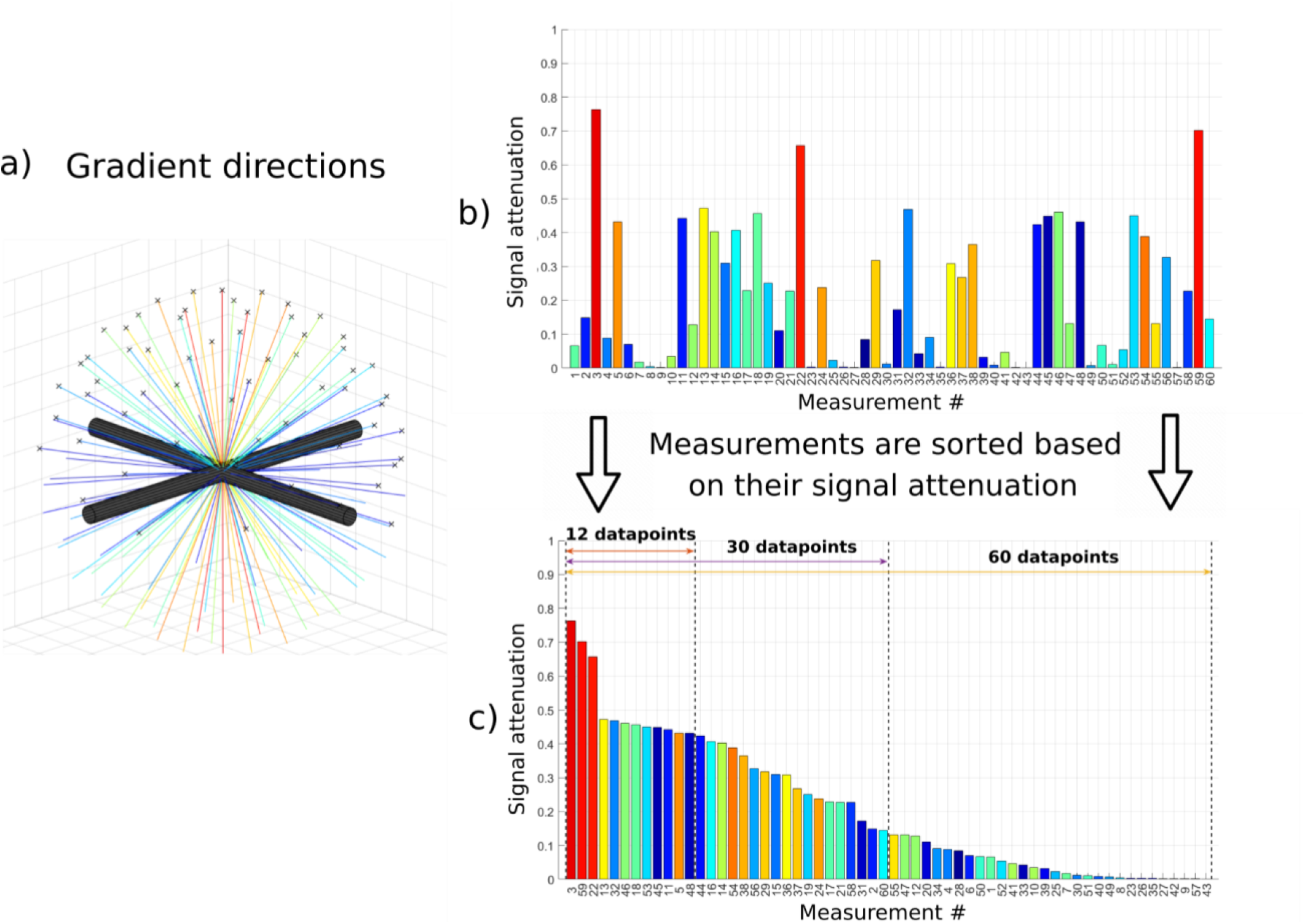
Effect of the gradient direction on the signal coming from two crossing fibre bundles. a) Gradient directions for an acquisition protocol with 60 measurements. b) Signal attenuation for each diffusion direction simulated for a substrate consisting of two perpendicular fibres / bundles each with Gamma diameter distributions of mean = 5.33 μm, std = 3.00 μm (the same as used in the ‘large axon’ simulations). c) Signal attenuations sorted in descending order and examples of subsets with different numbers of measurements.

In order to investigate whether this effect affects TDR, we divide the signal measurements into subsets and evaluate TDR for each different subset. The idea is to explore whether using a subset of gradient directions that creates the highest signal is more optimal (i.e. maximises TDR) compared to when using the full set of 60 uniformly distributed directions which includes both the high and the low signals. Note that the measurements are <u>already</u> acquired from all 60 uniformly distributed gradient directions, and this subset selection is happening at the post-processing stage, once the signal intensities have been ordered.

We use the definition of TDR outlined in Section 2.1. and calculate it for different subsets using the following equation:

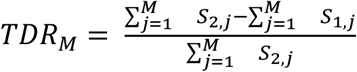

where M is the number of gradient directions in the subset (N corresponds to the full set of uniformly distributed directions and 1≤M≤N) and j=1..M is the index of the ordered signal measurements from the highest intensity to the lowest intensity (as shown in Figure 5c). The ordering is performed on the average of the two signals (*S*_1_ + *S*_2_)/2, in order to reduce the effect of noise on the sorting process. In our simulations we used HARDI acquisitions of 60 uniformly distributed gradient directions, hence N=60 and *TDR*_60_ is equivalent to the full original formulation of TDR (ie. signal S_1_ and S_2_ are averaged across all 60 gradient directions acquired). To illustrate this further: *TDR*_1_ is calculated using only one gradient direction - the one that provides maximum signal intensity; *TDR*_2_ uses the two strongest signal measurements to calculate the average, the one used in *TDR*_1_ and another one in the ordering etc.

The results for TDR calculated for every subset from *TDR*_1_ to *TDR*_60_ are provided in Figure 6 (a-d). Figure 6a shows the TDR values for a substrate consisting of one fibre bundle of parallel cylinders. Results are presented both for the noise-free scenario (blue circles) and the scenario where Rician noise (SNR = 20) has been added (orange error bars present the standard deviation over the noise instances). In the scenario without noise, TDR remains equal to the ground truth regardless of how many gradient directions are used in the TDR calculation. However, once simulated Rician noise is added to the signals, the results show a decrease in TDR and a reduction in the accuracy of the calculated TDR value as the number of gradient directions included in the calculation increases. In contrast, the precision of the calculation is improved with more measurements and hence for this substrate with one fibre bundle, using *TDR*_12_ which corresponds to the subset of ∼20% of ordered gradient directions that provide the highest signal values, yields an optimal balance between accuracy and precision of TDR estimates.

**Figure 6.**
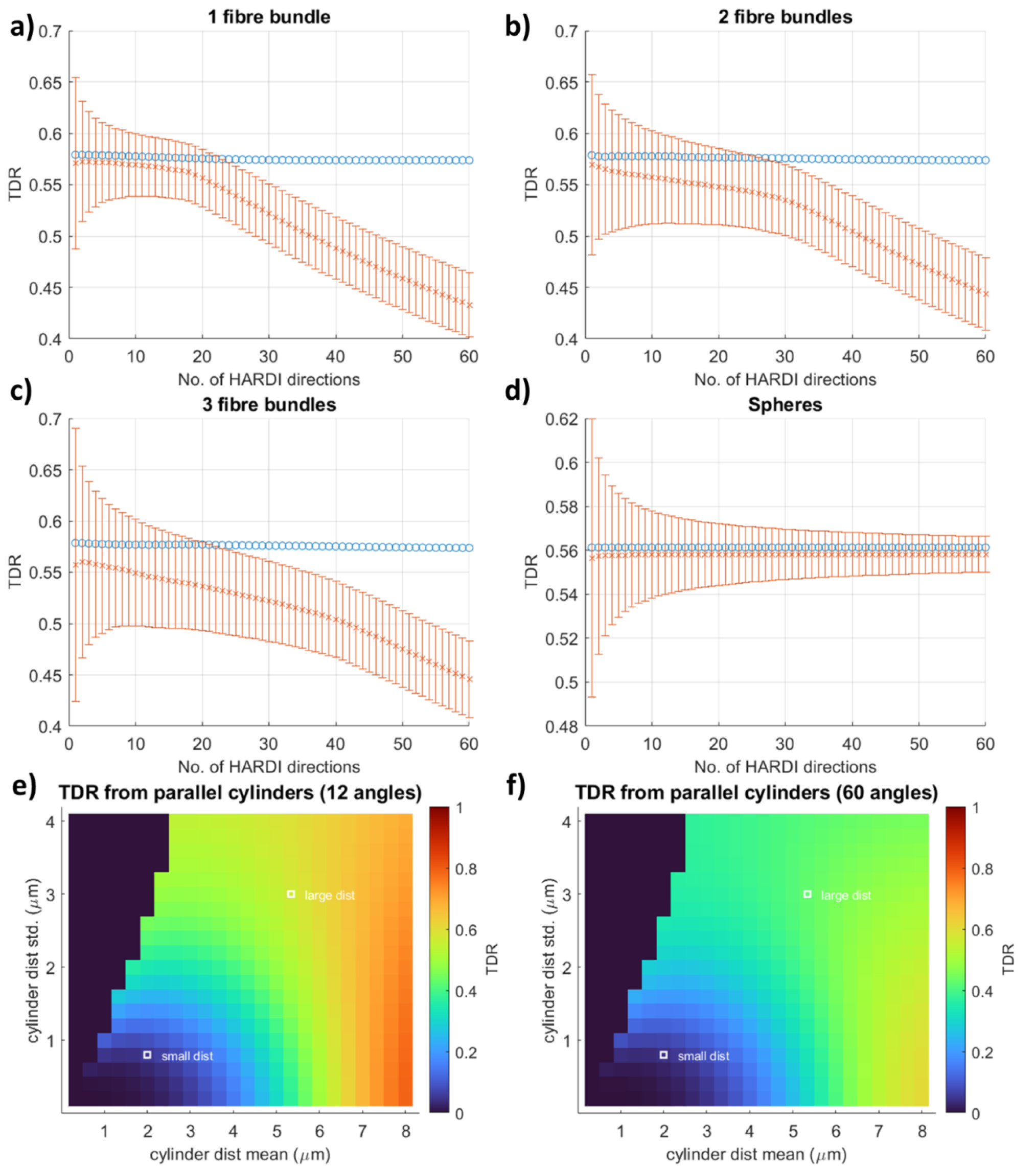
TDR values for noise free (blue circles, SNR = inf) and noisy (orange crosses, Rician noise with SNR = 20) signals as a function of the number of gradient directions included in the analysis in different substrates: a-c) bundles of cylinders with one, two or three fibre orientations crossing at 90°. All bundles have a Gamma distribution of sizes with mean = 5.33 μm and std =3.00 μm; d) a Gaussian distribution of spheres with mean = 7 μm and std = 0.5 μm. The orange error bars show 1 standard deviation of the estimated TDR over 100,000 noise instances. As illustrated in a-c), for the different fibre configurations, TDR values estimated from noisy data are below the expected noise-free values. All simulations are performed using the optimised sequence parameters for large cylinders.

Figures 6b and c show that similar effects occur for substrates consisting of two and three fibre bundles (fibre bundles are mutually perpendicular to allow for testing of the most extreme cases). The accuracy of estimated TDR lowers as the number of fibre bundles increases, however the trend that the accuracy reduces with increasing the size of the ordered subset is equally present suggesting that optimisation in the presence of Rician noise would be very beneficial and would provide higher TDR contrast.

For a substrate consisting of spheres (Figure 6d) the results are very different. Accuracy is pretty stable regardless of the number of signal measurements used for both the no-noise and noisy case. This is as expected since signal from spherical substrates is indifferent to gradient direction as opposed to the fibres which are highly sensitive to it. Similarly to the fibre simulations, the precision is improved with the number of measurements and hence the optimal solution for spherical substrates is that which maximises the number of measurements - i.e. the full set of HARDI acquisitions (*TDR*_60_ for our simulations). Hence, for isotropic pores, as well as uniformly oriented fibres (data not shown), using more gradient directions is optimal, with TDR values remaining accurate even when all directions are used.

We also note that the estimated TDR values for the substrates with one, two or three fibre bundles should ideally be the same, as the size distributions of the substrate cylinders are the same regardless of the fibre orientation distribution. When calculating the coefficient of variation of TDR values across the three substrates, for an SNR of 20 when 60 HARDI gradient directions have been acquired, we see a minimum for subsets with 27/60 gradient directions, as illustrated in Figure S3 in Supplementary Information. For other SNR levels and numbers of acquired gradient directions, the exact number of measurements in the optimal subsets might vary slightly, nevertheless, using a smaller subset of gradient directions appears optimal, also for SNR = 50 and 30 measured gradient directions, as illustrated in Figures S2-S3 in Supplementary Information. Overall, in all fibre scenarios considered, using ∼50% of the gradient directions acquired provides a more accurate TDR estimate than when all gradient directions are used; in the SNR 50 scenarios or when 60 HARDI gradient directions are acquired, using ∼33-50% of the directions also reduces the coefficient of variation between the three fibre scenarios compared to when all gradient directions are used.

Figures 6e and f show *TDR*_12_and *TDR*_60_, respectively, in the presence of noise, for the single fibre bundle scenario, across a wide range of different cylinder distributions. We can see that using all 60 directions results in the reduction of TDR compared to the ground truth (no-noise case, Figure 4a) for many large distributions: this also reduces the difference in TDR between large and small distributions, reducing the potential contrast of a TDR image. *TDR*_12_ is on the other hand considerably higher and closer to the ground truth: for example for the distribution of cylinders with a mean 5.33μm, *TDR*_12_is 1.6% lower compared to the ground truth while *TDR*_60_, is 24.5% lower - a greater than 15 times difference in percentage error - which in the real-world scenario could create contrast that is not sufficiently detectable.

### 3.2 Preclinical experiments

#### Optimised and non-optimised acquisition protocols

Diffusion MRI measurements for TDR contrast in ex-vivo spinal cord were acquired following the original, non-optimised, TDR approach, where the diffusion time is varied between the two shells, as well the optimised gradient waveforms proposed in this work. The measurements were repeated for a maximum gradient strength of 600 mT/m, a value widely available on pre-clinical systems as well as 2500 mT/m that is available on this particular scanner. The specific values for the non-optimised and optimised protocols are given below:

#### Non-optimised protocol

- Shell 1 consists of waveforms with short gradient duration and short diffusion time (G_max_= 600 mT/m: δ = 6.9 ms, Δ = 9 ms; G_max_= 2500 mT/m: δ = 2.2 ms, Δ = 4.5 ms). The optimal values from simulations were slightly adjusted to accommodate scanner constraints such as finite slew rates. This shell is used both for the original and the optimised TDR calculation, as prescribed by the simulation results.
- Shell 2n consists of waveforms with short gradient duration and long diffusion time (G_max_= 600 mT/m: δ = 6.9 ms, Δ = 34.6 ms; G_max_= 2500 mT/m: δ = 2.2 ms, Δ = 39 ms). The maximum diffusion time given a fixed Δ+δ value was chosen. TDR values computed from Shell 1 and Shell 2n are referred to as non-optimised.

#### Optimised protocol

- Shell 1 is the same as for the non-optimised protocol.
- Shell 2o consists of waveforms with long gradient duration and long diffusion time (G_max_= 600 mT/m: δ = 14 ms, Δ = 27.5 ms; G_max_= 2500 mT/m: δ = 15 ms, Δ = 26.5 ms). The timing parameters are adapted from the numerical optimization. TDR values computed from Shell 1 and Shell 2o are referred to as optimised.

The signal attenuation profiles for these gradient waveforms plotted against axon diameter are presented in Figure 7, illustrating that indeed the difference between Shell 2o and Shell 1 (optimised) is larger than between Shell 2n and Shell 1 (not-optimised).

**Figure 7.**
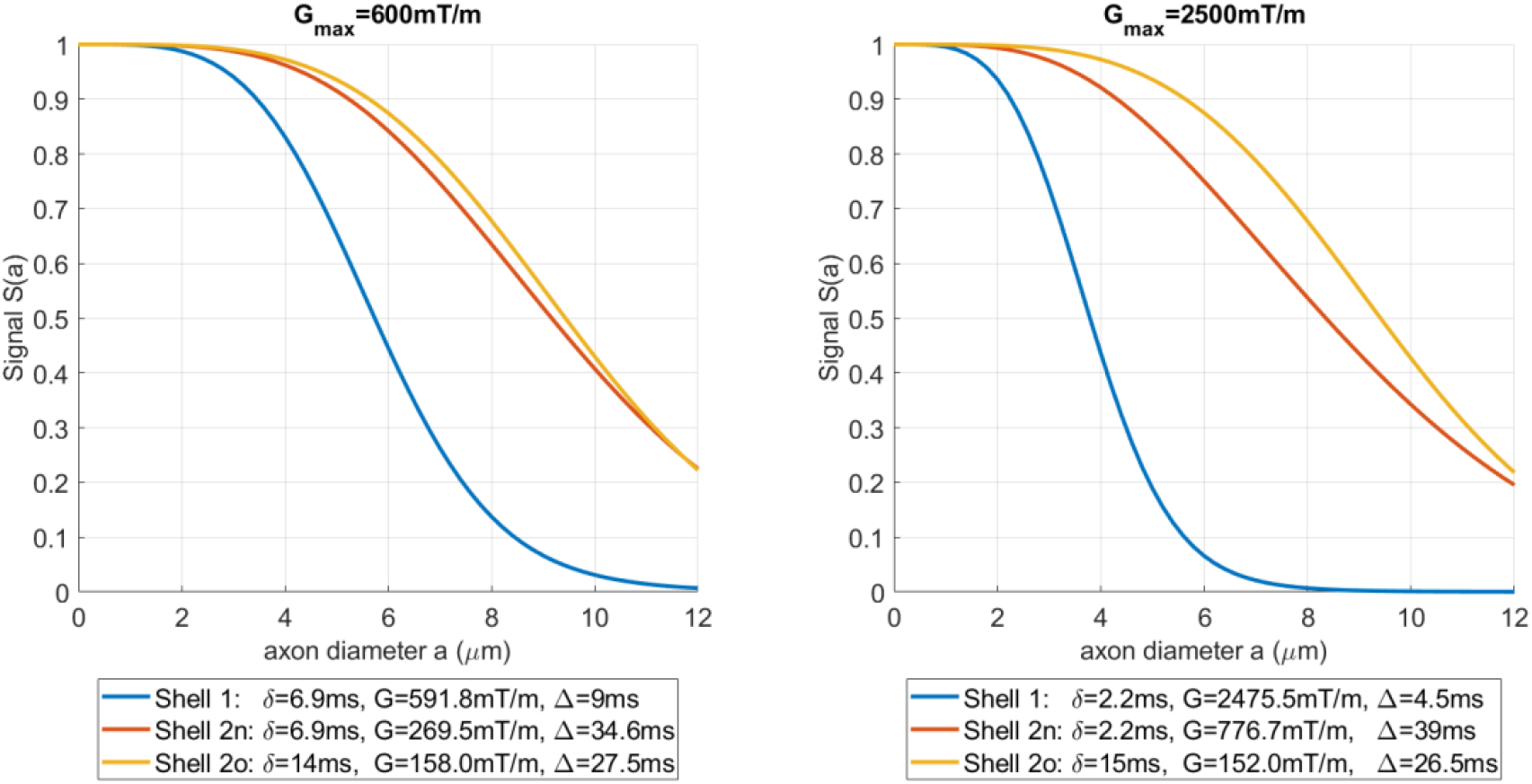
Simulations for signal attenuation profiles as a function of cylinder diameter for the three shells used in the pre-clinical acquisition, both for G_max_= 600 mT/m (left) and G_max_= 2500 mT/m. The signals were simulated for single cylinders and diffusion gradients perpendicular to the fibre with intrinsic diffusivity of D = 2 μm^2^/ms.

#### TDR contrast in spinal cord

Figure 8a illustrates the data acquired in the ex-vivo rat spinal cord, where WM and GM regions are outlined on the b0 image. The normalised diffusion weighted maps are shown for the three different waveforms when the gradient is either close to parallel or perpendicular to the spinal cord fibres. Indeed, there is a pronounced change in contrast between the different shells. For the perpendicular direction, the signal in white matter increases between shell 1 and shells 2n & 2o, while the signal in gray matter decreases. For the parallel direction, the change is less pronounced in white matter, nevertheless there is still a pronounced decrease in gray matter.

**Figure 8.**
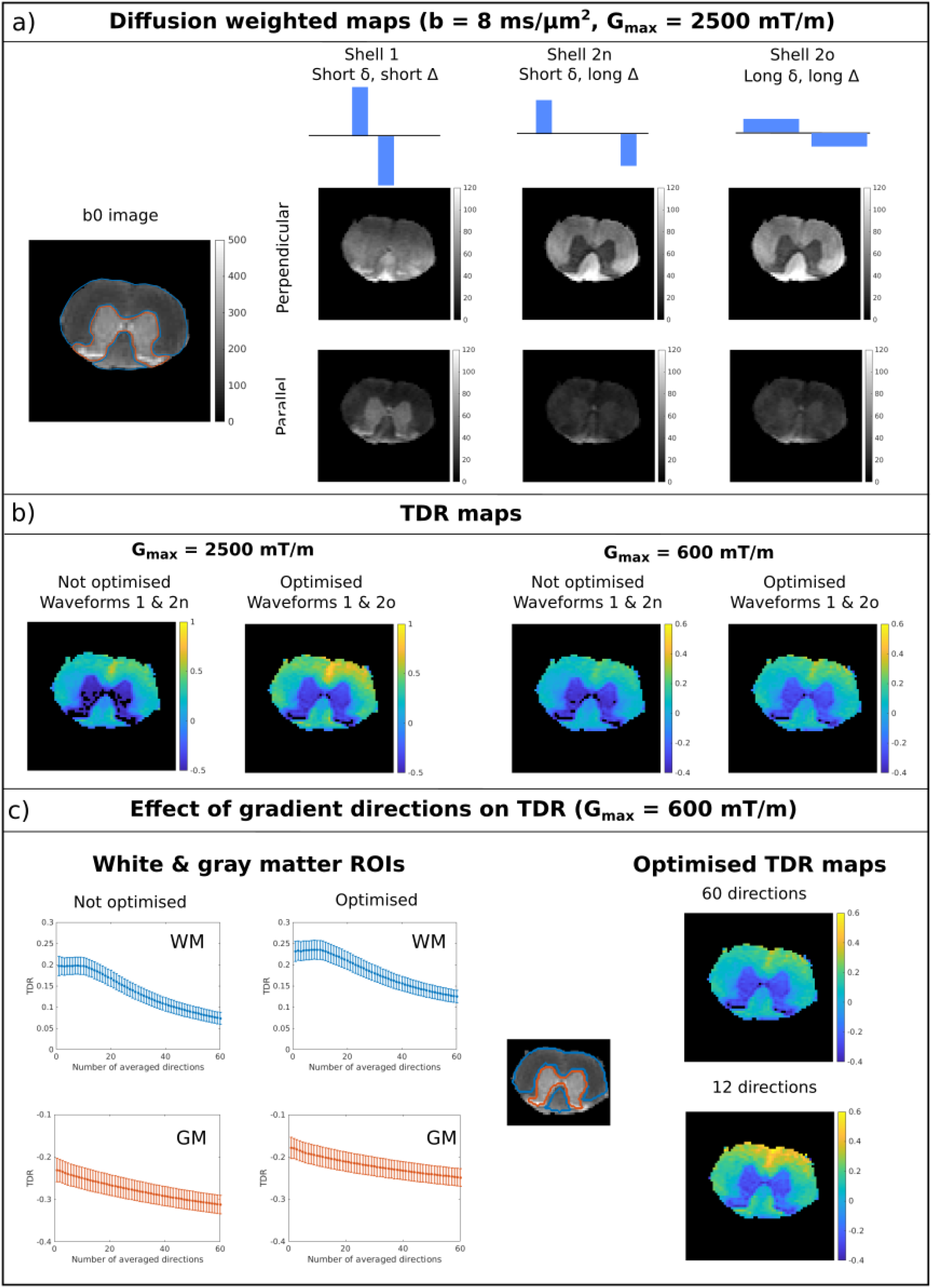
Optimisation of TDR acquisition including waveforms (a-b) and number of gradient directions (b). a) left: T2 weighted image of the rat spinal cord without diffusion gradients. White matter and gray matter regions are delineated with blue and orange contours, respectively. Right: schematic depiction of the gradient waveforms for the three shells (top) and the corresponding diffusion weighted maps for a perpendicular (middle) and a parallel (bottom) gradient direction. b) TDR maps for maximum gradient strength of 600 and 2500 mT/m for optimised and non-optimised gradient waveforms. c) left: TDR values as a function of how many gradient directions were included in the signal average for white matter (blue) and gray matter (orange) ROIs. right: TDR maps for measurement subsets with 12/60 and 60/60 directions.

Figure 8b presents TDR maps calculated based on all gradient directions for the two scenarios with G_max_= 600 mT/m as well as for G_max_= 2500 mT/m. In spinal cord white matter, TDR contrast is positive. For G_max_= 600 mT/m, optimised TDR values in WM are between 0 and 0.4, matching a wide range of axonal distributions presented in Figure 4a for the same gradient strength. On the other hand, in gray matter, TDR values are negative, showing that a model of pure restriction (either in cylinders and/or spheres) does not accurately represent the diffusion time dependence in the tissue.

For both gradient strengths, we see that the TDR contrast provided by the optimised pair of gradient waveforms (i.e. Shell 1 and Shell 2o) is higher than the values from the non-optimised pair (Shell 1 and Shell 2n). Moreover, TDR values obtained with stronger gradients are higher, matching the simulation results presented in Supplementary Information in Figure S1, as well as previous results regarding estimating axon diameters ^26,33,37^.

To assess the effect of gradient direction on the TDR estimate in the spinal cord, where white matter tracts are highly anisotropic, the gradient directions were voxelwise sorted based on the average signal intensity in the three shells. Then, TDR was computed from subsets of gradient orientations with an increasing number of directions, as detailed in Figure 5.

Figure 8c illustrates the effect of using a subset of gradient directions in the computation of TDR. For WM, we see that both the optimised and non-optimised TDR values decrease as more gradient directions are included in the signal average. The plots also reflect that including aprox 20% of the data points (i.e. 12/60 directions) provides a good balance towards maximising TDR while minimising the effect of noise, corroborating the simulation results. In GM, there is also a dependence of TDR on the number of measurements, likely due to the directionality of the neurites^74^. Similar results are observed for the second spinal cord segment.

Overall, for a given gradient strength and echo time, the highest TDR contrast in spinal cord white matter is obtained using the optimised pair of gradient waveforms and approx 12/60 directions from a fully acquired HARDI shell.

Figure 9 compares TDR with axon diameter values reported in the literature in 6 different WM ROIs. The TDR values are calculated voxelwise from the optimised waveforms using a subset of 12 gradient directions, to maximise the contrast. Then, the mean TDR value is each ROI is computed. Strong correlations between mean TDR values and axon diameter are observed, both for G_max_= 600 mT/m, with a correlation coefficient of 0.86 (p<0.01), as well as for G_max_= 2500 mT/m with a correlation coefficient of 0.89 (p<0.01). Moreover, the TDR values for the two spinal cord segments, that were mounted and imaged separately, are highly consistent.

**Figure 9.**
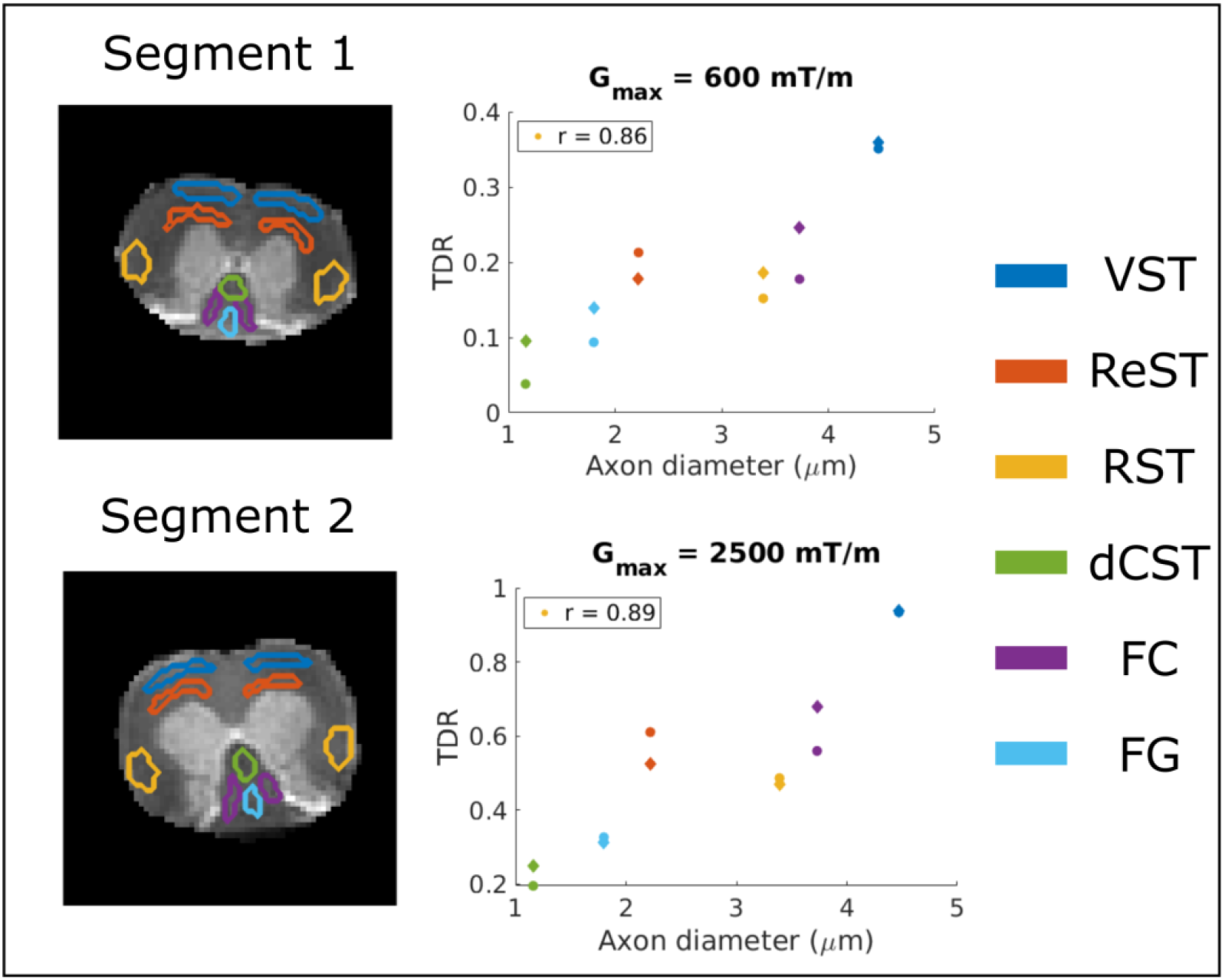
Comparison between TDR values and axon diameters reported in the literature in 6 spinal cord WM ROIs (Vestibulospinal (VST) - blue, Reticulospinal (ReST) - orange, Rubrospinal (RST) - yellow, dorsal corticospinal (dCST) - green, Funiculus Cuneatus (FC) - purple and Funiculus Gracilis (FG) - light blue). The ROIs are manually delineated and colour coded as illustrated on the left. Each segment is represented with a different marker shape in the correlation plot. Strong correlations between TDR and axon diameter are observed, both for G_max_= 600 mT/m (correlation coefficient r = 0.86) and G_max_= 2500 mT/m (correlation coefficient r = 0.89).

## 4. Discussion

This work employs numerical simulations to optimise the dMRI acquisition for maximal TDR contrast. The simulation results are validated by ex-vivo diffusion MRI data from both WM and GM tissues in rat spinal cord.

### TDR contrast and waveform optimisation

Through our investigation of pulse sequence shapes for standard single diffusion encoding experiments, we find that optimised TDR requires short-δ, high-G sequences contrasted with long-δ, low-G sequences. These results are consistent for a wide range of substrates (e.g. restricted diffusion in cylinders and spheres with different size distributions) and sequence constraints (maximum gradient strength, total gradient duration, etc), and are in line with the optimisation results of previous studies^18,33,37,40^.

The contrast from the optimised TDR is sensitive to a range of restriction sizes, as illustrated in Figure 4, and has the potential to distinguish distributions containing large axons from those containing small axons. As expected, size distributions with different combinations of mean and standard deviation can yield very similar TDR values, as illustrated by areas with similar colours in Figure 4. This effect is more pronounced for distributions with longer tails, such as the gamma distribution used for cylindrical substrates, compared to normal distributions used for spherical substrates, where we observe a better correspondence between the mean of the distribution and TDR values. This known tail-weighting effect is due to the fact that the signal contribution of each cylinder to the total measured signal is volume-weighted^18,20^. As reference, for axon diameter distributions typically found in human brain tissue (i.e. < 5 μm), TDR contrast is on the order of 10% or less^55^.

As illustrated both through simulations and experiments, the TDR contrast depends on the maximum gradient strength available on the scanner, and is sensitive to axon diameters that are above the resolution limit, i.e. the minimum diameter that can be distinguished^18,33,37^. For sizes below the resolution limit, TDR is indistinguishable from zero.

### Effect of extracellular space

The TDR exploits ultra-high b-values to suppress the contribution from water diffusing in the extracellular space and enhance the sensitivity of the dMRI measurements to water diffusing in the restricted intracellular space. In this study we choose to investigate sequences with b = 8000 s/mm^2^, as it is large enough to attenuate the signal from extracellular space, yet can be achieved on pre-clinical and high performance clinical scanners for a range of gradient waveforms, as required in TDR. Regarding the signal attenuation in the extracellular space, we used the simulation results presented in Figure 2 from a previous work by Grussu et al.^24^, that consider Gamma distributions of cylinders with a realistic packing, to estimate the relative contribution of the extracellular space. Thus, for the substrates with small axons (mean diameter 3 μm), close to the mean axon diameter across the different ROIs in the rat spinal cord, the relative signal contribution of the extracellular space is less than 5% using the diffusivity and kurtosis values at 30 ms. For spinal cord ROIs with larger diameters (up to 4.5 μm) this contribution might increase, nevertheless, it will be significantly less than 20%, the value calculated for large axons (mean diameter 7 μm) with high packing density.

### TDR in the presence of noise

In the original TDR, it was proposed to use the direction-averaged signal to remove/mitigate the bias due to orientation dispersion. In this work we show that, when imaging structures consisting of one or several anisotropic fibre bundles (e.g. coherent white matter fibres, but also regions with crossing fibres), in the presence of Rician noise, the TDR contrast can be further improved by using only a subset of the highest-signal gradient directions. This happens because for some gradient directions the measured signal is very low and decays to the noise floor, therefore adding a bias in the powder averaged signal and subsequent TDR calculation. Thus, removing these measurements improves the accuracy of TDR estimation. For relatively coherent fibres, both simulations and experimental results suggest that using approx 20% of the measurements is optimal. For more complex fibre configurations (e.g. two or three crossing fibre bundles), larger subsets are more appropriate, with the exact optimal percentage of gradient directions depending on the SNR as well as the fibre configurations. If we are to choose a single percentage, for example to calculate TDR across the brain where there are voxels with different fibre orientations, our simulations suggest that considering a subset with e.g. ∼50% of measurements, improves TDR accuracy and decreases its variance across fibre orientations compared to using the entire dataset. Nevertheless, the optimal percentage should be decided based on the SNR of the data, the number of HARDI directions and expected fibre orientations in the sample.

This bias in TDR can also be mitigated by employing real valued data instead of magnitude data and removing the effect of the Rician noise floor. Indeed, simulations employing Gaussian noise result in similar TDR values for different numbers of gradient directions (data not shown).

### TDR in ex-vivo spinal cord: white matter

The TDR contrast in the ex-vivo rat spinal cord closely follows the predictions from simulations, with higher values for the optimised waveforms and ∼20% of the directions. We have also observed very strong correlations between TDR and axon diameter values reported in the literature in different ROIs, with correlation coefficients above 0.85 both for a weaker gradient (600 mT/m) and a stronger gradient (2500 mT/m). As discussed above, TDR is sensitive to the volume weighted distribution of axons, and therefore the mean diameter calculated from electron microscopy might not be the best quantity for comparison, nevertheless, there is a clear trend, as illustrated in Figure 9. The trends of TDR are also consistent with previous MRI correlates of axon diameter, both using diffusion data as well as relaxometry ^30,75–77^.

### TDR in ex vivo spinal cord: gray matter

In *ex vivo* spinal cord gray matter we measure negative TDR values. These results are not consistent with simulations of TDR when diffusion is totally restricted in spheres and/or cylinders, suggesting that effects, other than pure restriction, influence the diffusion time dependence in gray matter. These results show for the first time that for measurements with high diffusion weighting (i.e. 8000 s/mm^2^), the signal is showing a pronounced decrease with increasing diffusion time in spinal cord gray matter. This is consistent with recent preclinical studies focusing on brain gray matter, suggesting that inter-compartmental exchange plays an important role on the diffusion time dependence, both *in vivo* and *ex vivo*^*43,44,78*^. In this case, TDR may reflect exchange between neurites and extracellular space, between soma and extracellular space and between soma and neurites^40,43,44^. Since TDR was originally proposed to map restriction effects in the WM, here we focus our optimization and investigation on WM. The observed discrepancy between simulated and measured TDR values in the GM are of great interest, but require further dedicated investigations, which goes beyond the scope of this work.

### Going beyond single diffusion encoding

The TDR contrast could be further improved by including other waveforms in the optimization, for example oscillating gradients^33^, and a similar approach has been explored recently in simulations using such waveforms^19^. If high frequency oscillating gradients cannot usually achieve the desired high b-values for a practical echo time, low frequency oscillations could potentially improve TDR contrast, as they were shown in the past to increase the sensitivity towards small axon diameters. Nevertheless, these sequences are not widely available, neither on clinical nor on pre-clinical scanners, therefore here we focused on standard single diffusion encoding acquisitions. For larger restriction sizes, using SDE sequences in a stimulated echo sequence rather than the standard spin-echo preparation might be beneficial as it allows to reach larger gradient separations, a period where the signal is governed by a slower T1 decay rather than the faster T2 decay.

### Limitations

In this work only simple and ideal geometrical substrates (parallel cylinders and spheres) are considered with no exchange between compartments, so a full exploration of negative TDR values is not possible. The inclusion of more complex substrate models potentially involving dispersion, axon curvature^79,80^, beads^81,82^, and exchange might provide more insight into the behaviour of TDR across a wider range of possible microstructures. Another simulation limitation is that noise is simulated using the Rician distribution; strictly speaking, noise across a multi-channel receive coil (e.g, 32 channels for many clinical scanners) should have a non-central Chi distribution. Moreover, TDR contrast could be further improved by considering other gradient shapes, such as oscillating gradients or spin preparations, e.g. stimulated echo, nevertheless, this would require the use of new sequences that are not widely available at the moment.

TDR requires special care when interpreting the results. As TDR is a single-dimensional measure of restricted space, some structures with different compositions will return identical TDR values. Examples of these structures can be determined through examination of the isocolours in Figure 4, where cylinder and sphere distributions with a range of means and standard deviations can return the same TDR values (and hence are coloured the same colour in Figure 4). More generally, when moving beyond substrates consisting simply of cylinders or spheres, there will be a range of microstructure compositions which we can expect to return similar values of TDR. This highlights the function of TDR as a method for mapping areas containing large restrictions rather than as a method for directly determining pore diameters.

TDR is also affected by the inherent sensitivity of the acquisition and the diffusion signal to the pore sizes. As shown in previous work^33,37^, the sensitivity of the diffusion signal to axon diameter is very dependent on the gradient strength and SNR. These can be used to calculate a size resolution limit below which the signal has minimal sensitivity. Thus, the simulations employed in this study cover the settings of high-performance gradient systems (i.e. the lowest gradient strength we consider in this paper is 300mT/m), where the TDR contrast can provide sensitivity to axon diameters. Further work will also assess gradient amplitudes more commonly available on clinical scanners (e.g. G < 100 mT/m) to map larger restriction sizes that can be found in different tissue types, for instance in tumours. While its dependence on the resolution limit is very similar to model-based approaches, TDR has an advantage that it does not try to fit but rather just map areas where the voxels contain pores of sizes above the resolution limit. Hence, its robustness, sensitivity and interpretation will be much improved and more appropriate for some applications compared to the model-based approaches.

## 5. Conclusion

This work employs simulations and pre-clinical data to show the potential of TDR contrast to characterise restricted diffusion for a wide range of tissue microstructures featuring cylindrical and/or spherical size distributions. Importantly, the TDR contrast can be enhanced by using optimised gradient waveforms, contrasting a short δ + high G pulse and a long δ + low G pulse, as well as a subset of gradient directions from the acquired shells. In ex-vivo experiments on rat spinal cord, TDR can successfully characterise spinal cord white matter microstructure, showing a strong correlation with axon diameter values from quantitative histology. Overall these results show that the recently proposed TDR approach has a great potential and is a very promising alternative (or potentially complement) to model-based approaches for mapping pore sizes in tissue.

## Acknowledgements

This research was funded in part by the ESPRC (grant reference EP/R513143/1).

The authors acknowledge the vivarium of the Champalimaud Center for the Unknown, a facility of CONGENTO financed by Lisboa Regional Operational Programme (Lisboa 2020), project LISBOA01-0145-FEDER-022170. AI and RC are supported by “la Caixa” Foundation (ID 100010434) and from the European Union’s Horizon 2020 research and innovation programme under the Marie Skłodowska-Curie grant agreement No 847648, fellowship code CF/BQ/PI20/11760029. NS is supported by the European Research Council (ERC) under the European Union’s Horizon 2020 research and innovation programme (Starting Grant, agreement No. 679058).

MP is supported by the UKRI Future Leaders Fellowship MR/T020296/2

This research was funded in part by a Wellcome Trust Investigator Award (096646/Z/11/Z) and a Wellcome Trust Strategic Award (104943/Z/14/Z). For the purpose of open access, the author has applied a CC BY public copyright licence to any Author Accepted Manuscript version arising from this submission.

## Supplementary Information

As well as our optimisations in Simulation 1, which use hardware constraints corresponding to a typical pre-clinical scanner, we additionally optimise TDR for G < 2700 mT/m, Δ+δ < 45 ms corresponding to a high-gradient pre-clinical system and G < 300 mT/m, Δ+δ < 80 ms corresponding to the Connectom scanner. We have also included typical duration for the refocusing pulse between the diffusion gradients, leading to Δ-δ > 2ms for the preclinical settings and Δ-δ > 7ms for the Connectom settings. Results when optimising TDR for a substrate consisting of large parallel cylinders (as specified in Figure 2) are displayed in Figure S1.

Overall, when optimising TDR for a large cylinder substrate for our three different sets of scanner constraints, we find the following parameter sets:

- G<300mT/m, Δ+δ < 80ms: S_1_ : Δ=16.62 ms, δ=9.62 ms; S_2o_: Δ=53.58 ms, δ=26.42 ms
- G<600mT/m, Δ+δ < 45ms: S_1_ : Δ=8.87 ms, δ=6.87 ms; S_2o_: Δ=30.92 ms, δ=14.08 ms
- G<2700mT/m, Δ+δ < 45ms: S_1_ : Δ=4.12ms, δ=2.12 ms; S_2o_: Δ=30.92 ms, δ=14.08 ms

**Figure S1:**
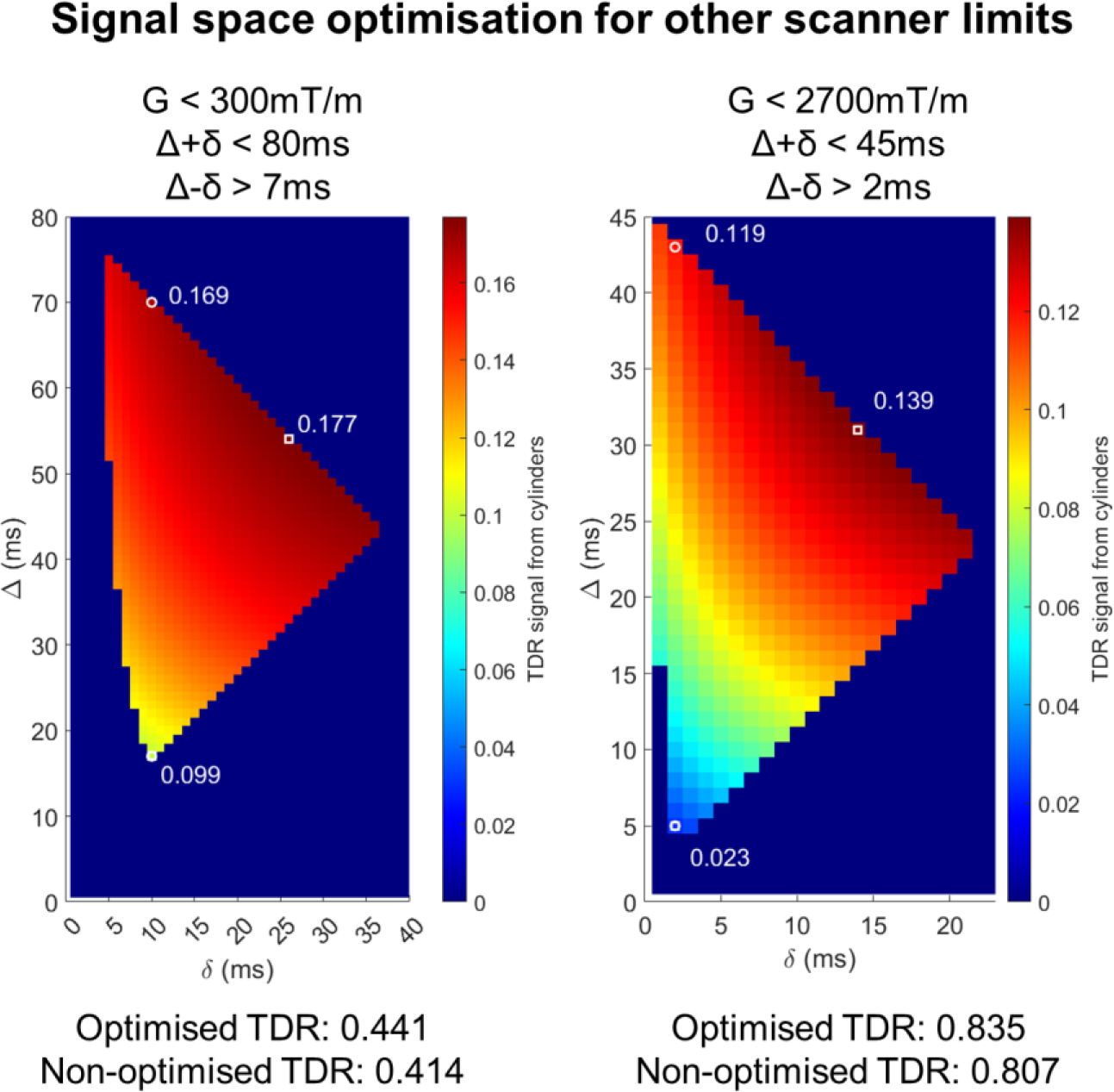
Optimisation results maximising TDR for other scanner constraints. Maps show the diffusion weighted signal for sequences with b=8000 s/mm^2^and various δ/Δ combinations for the ‘large axon’ substrate illustrated in Figure 2. Signal values are averaged over 60 directions as in the original formulation of TDR. White markers indicate the non-optimised (circle) and optimised (square) sequences, respectively. Optimal parameters found through nonlinear optimisation are: [G<300mT/m: S_1_ : Δ=16.62 ms, δ=9.62 ms; S_2o_: Δ=53.58 ms, δ=26.42 ms] and [G<2700mT/m: S_1_ : Δ=4.12ms, δ=2.12 ms; S_2o_: Δ=30.92 ms, δ=14.08 ms].

As discussed in the results of Simulation 3, calculating TDR using a subset of the acquired gradient directions that provide the highest signals, can increase the accuracy of the TDR estimate in the presence of Rician noise. For the same substrates presented in Figure 6, here we explore this effect for other acquisition scenarios, with different numbers of gradient directions (30 instead of 60) and SNR levels (50 instead of 20).

We find that in all scenarios, calculating TDR using subsets consisting of fewer gradient directions improves the accuracy of TDR (Figure S2).

We also consider the coefficient of variation of the mean TDR in the presence of noise, calculating the coefficient of variation across the three different fibre scenarios. As the three fibre scenarios all include fibres bundles with identical diameter distributions, ideally TDR should return the same value for all three scenarios, and the coefficient of variation should be zero.

We find that when SNR = 50, the coefficient of variation is minimised by using ∼33% of the available gradient directions (21/60 and 10/30 directions respectively). Alongside our results for SNR = 50 in Figure S2, this suggests that calculating TDR using a subset of the available directions is optimal.

When SNR = 20 and 60 gradient directions are acquired in total, using 27/60 or 45% is a clear local minimum in the coefficient of variation plot; as the true minimum is found at 2/60 directions, where the variation in calculated TDR due to noise is unacceptably large (Figure 6 a-d) this also suggests using roughly half the gradient directions available is a suitable choice for calculating TDR in the presence of noise.

**Figure S2:**
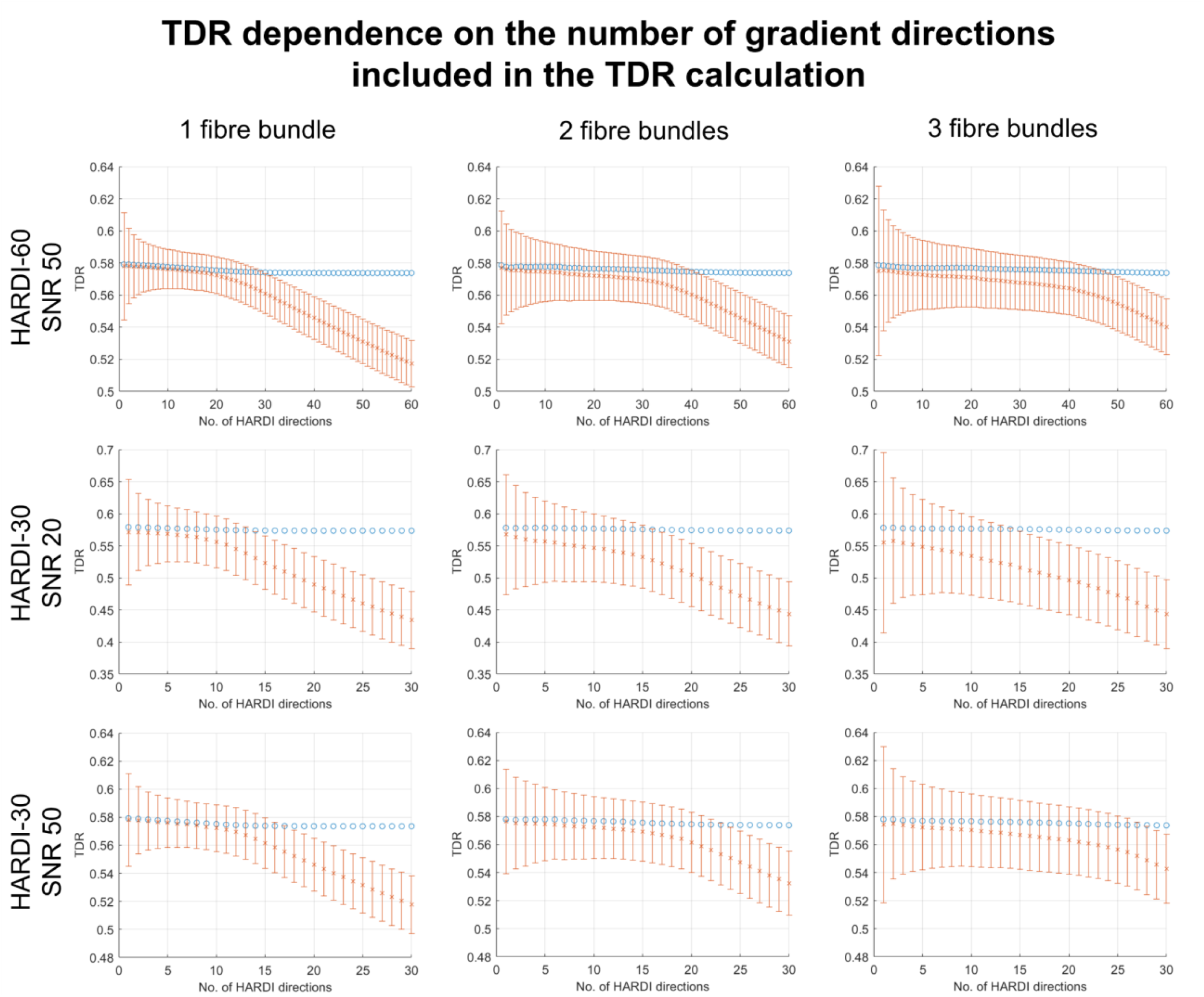
TDR values for noise free (blue circles, SNR = inf) and noisy (orange crosses, Rician noise with SNR as shown) signals as a function of the number of gradient directions included in the analysis, in different substrates consisting of bundles of cylinders with one, two or three fibre orientations crossing at 90°. All bundles have a Gamma distribution of sizes with mean = 5.33 μm and std =3.00 μm. The orange error bars show 1 standard deviation of the estimated TDR over 100,000 noise instances. All data are acquired using the optimised sequence parameters for large cylinders.

**Figure S3:**
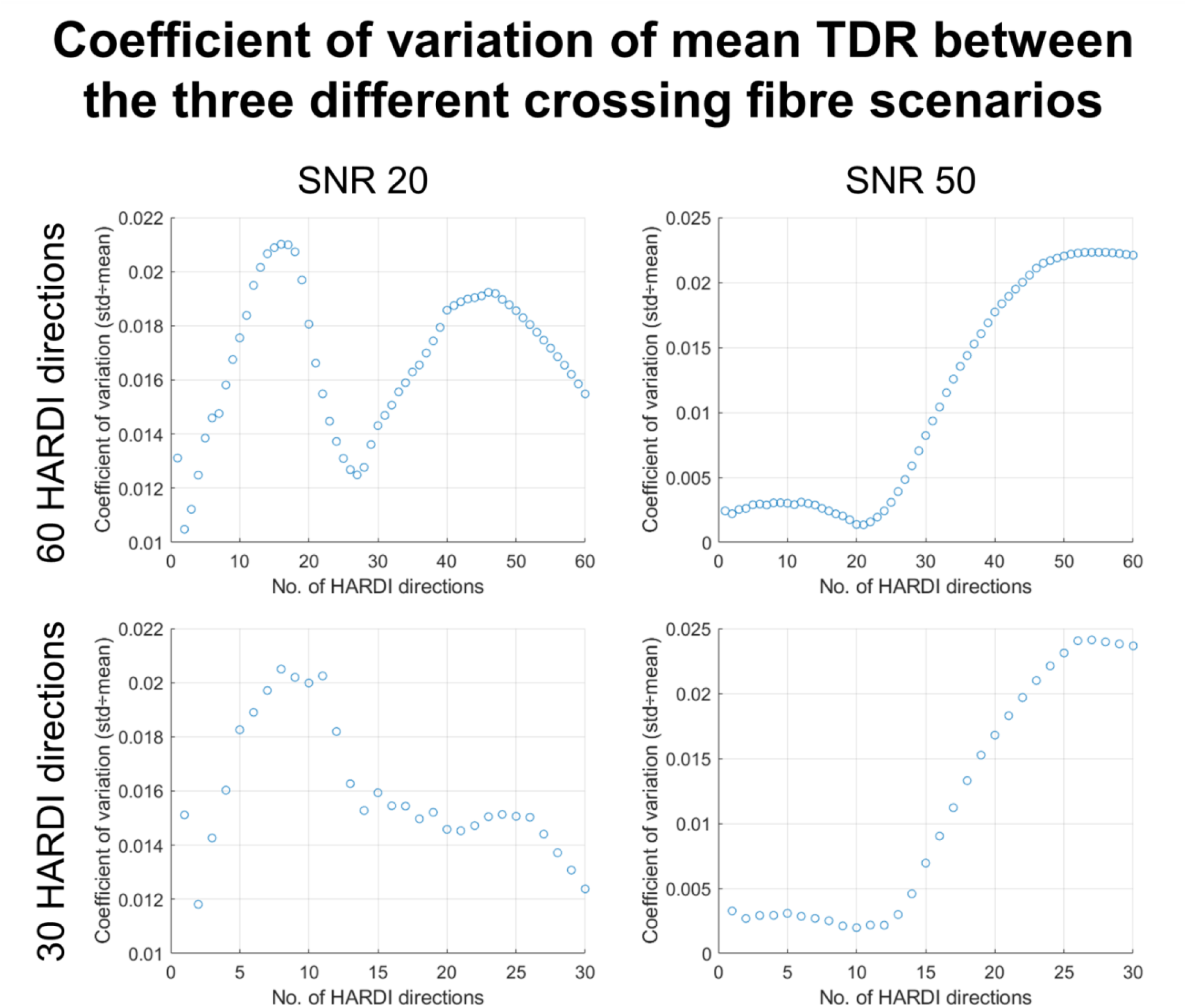
Coefficient of variation of mean TDR between the three scenarios with 1,2 and 3 fibre directions. In simulations with multiple fibre directions the directions chosen are perpendicular. Simulations are run four times, at SNR = 20 and SNR 50 for HARDI shells consisting of 60 and 30 gradient directions.

## References

1. Lawrence, K. E. et al. Age and sex effects on advanced white matter microstructure measures in 15,628 older adults: A UK biobank study. Brain Imaging Behav. 15, 2813–2823 (2021).

2. Tian, L. & Ma, L. Microstructural Changes of the Human Brain from Early to Mid-Adulthood. Front. Hum. Neurosci. 11, 393 (2017).

3. Dubois, J. et al. The early development of brain white matter: a review of imaging studies in fetuses, newborns and infants. Neuroscience 276, 48–71 (2014).

4. Sexton, C. E. et al. Accelerated changes in white matter microstructure during aging: a longitudinal diffusion tensor imaging study. J. Neurosci. 34, 15425–15436 (2014).

5. Lebel, C., Caverhill-Godkewitsch, S. & Beaulieu, C. Age-related regional variations of the corpus callosum identified by diffusion tensor tractography. Neuroimage 52, 20–31 (2010).

6. Hursh, J. B. CONDUCTION VELOCITY AND DIAMETER OF NERVE FIBERS. American Journal of Physiology-Legacy Content 127, 131–139 (1939).

7. Ritchie, J. M. On the relation between fibre diameter and conduction velocity in myelinated nerve fibres. Proceedings of the Royal Society of London. Series B. Biological Sciences 217, 29–35 (1982).

8. Heads, T., Pollock, M., Robertson, A., Sutherland, W. H. & Allpress, S. Sensory nerve pathology in amyotrophic lateral sclerosis. Acta Neuropathol. 82, 316–320 (1991).

9. Cluskey, S. & Ramsden, D. B. Mechanisms of neurodegeneration in amyotrophic lateral sclerosis. Mol. Pathol. 54, 386–392 (2001).

10. Bezchlibnyk, Y. B. et al. Neuron somal size is decreased in the lateral amygdalar nucleus of subjects with bipolar disorder. J. Psychiatry Neurosci. 32, 203–210 (2007).

11. Dukkipati, S. S., Garrett, T. L. & Elbasiouny, S. M. The vulnerability of spinal motoneurons and soma size plasticity in a mouse model of amyotrophic lateral sclerosis. J. Physiol. 596, 1723–1745 (2018).

12. Taylor, D. C., Falconer, M. A., Bruton, C. J. & Corsellis, J. A. Focal dysplasia of the cerebral cortex in epilepsy. J. Neurol. Neurosurg. Psychiatry 34, 369–387 (1971).

13. Baliyan, V., Das, C. J., Sharma, R. & Gupta, A. K. Diffusion weighted imaging: Technique and applications. World J. Radiol. 8, 785–798 (2016).

14. Drake-Pérez, M., Boto, J., Fitsiori, A., Lovblad, K. & Vargas, M. I. Clinical applications of diffusion weighted imaging in neuroradiology. Insights Imaging 9, 535–547 (2018).

15. Le Bihan, D. & Iima, M. Diffusion Magnetic Resonance Imaging: What Water Tells Us about Biological Tissues. PLoS Biol. 13, e1002203 (2015).

16. Stanisz, G. J., Szafer, A., Wright, G. A. & Henkelman, R. M. An analytical model of restricted diffusion in bovine optic nerve. Magn. Reson. Med. 37, 103–111 (1997).

17. Assaf, Y., Blumenfeld-Katzir, T., Yovel, Y. & Basser, P. J. AxCaliber: a method for measuring axon diameter distribution from diffusion MRI. Magn. Reson. Med. 59, 1347–1354 (2008).

18. Alexander, D. C. et al. Orientationally invariant indices of axon diameter and density from diffusion MRI. Neuroimage 52, 1374–1389 (2010).

19. Harkins, K. D., Beaulieu, C., Xu, J., Gore, J. C. & Does, M. D. A simple estimate of axon size with diffusion MRI. Neuroimage 227, 117619 (2021).

20. Veraart, J. et al. Noninvasive quantification of axon radii using diffusion MRI. Elife 9, e49855 (2020).

21. Kakkar, L. S. et al. Low frequency oscillating gradient spin-echo sequences improve sensitivity to axon diameter: An experimental study in viable nerve tissue. Neuroimage 182, 314–328 (2018).

22. Huang, S. Y. et al. High-gradient diffusion MRI reveals distinct estimates of axon diameter index within different white matter tracts in the in vivo human brain. Brain Struct. Funct. 225, 1277–1291 (2020).

23. Duval, T. et al. In vivo mapping of human spinal cord microstructure at 300mT/m. Neuroimage 118, 494–507 (2015).

24. Grussu, F. et al. Relevance of time-dependence for clinically viable diffusion imaging of the spinal cord. Magn. Reson. Med. 81, 1247–1264 (2019).

25. Sepehrband, F., Alexander, D. C., Kurniawan, N. D., Reutens, D. C. & Yang, Z. Towards higher sensitivity and stability of axon diameter estimation with diffusion-weighted MRI. NMR Biomed. 29, 293–308 (2016).

26. Fan, Q. et al. Axon diameter index estimation independent of fiber orientation distribution using high-gradient diffusion MRI. Neuroimage 222, 117197 (2020).

27. Barazany, D., Basser, P. J. & Assaf, Y. In vivo measurement of axon diameter distribution in the corpus callosum of rat brain. Brain 132, 1210–1220 (04 2009).

28. Zhang, H., Dyrby, T. B. & Alexander, D. C. Axon diameter mapping in crossing fibers with diffusion MRI. Med. Image Comput. Comput. Assist. Interv. 14, 82–89 (2011).

29. Ong, H. H. & Wehrli, F. W. Quantifying axon diameter and intra-cellular volume fraction in excised mouse spinal cord with q-space imaging. Neuroimage 51, 1360–1366 (2010).

30. Shemesh, N. Axon Diameters and Myelin Content Modulate Microscopic Fractional Anisotropy at Short Diffusion Times in Fixed Rat Spinal Cord. Frontiers in Physics 6, (2018).

31. Xu, J. et al. Mapping mean axon diameter and axonal volume fraction by MRI using temporal diffusion spectroscopy. Neuroimage 103, 10–19 (2014).

32. Bar-Shir, A. & Cohen, Y. High b-value q-space diffusion MRS of nerves: structural information and comparison with histological evidence. NMR Biomed. 21, 165–174 (2008).

33. Drobnjak, I., Zhang, H., Ianuş, A., Kaden, E. & Alexander, D. C. PGSE, OGSE, and sensitivity to axon diameter in diffusion MRI: Insight from a simulation study. Magn. Reson. Med. 75, 688–700 (2016).

34. Siow et al. Axon radius estimation with Oscillating Gradient Spin Echo (OGSE) Diffusion MRI. Diffusion Fundamentals 18, 1–6 (2013).

35. Does, M. D., Parsons, E. C. & Gore, J. C. Oscillating gradient measurements of water diffusion in normal and globally ischemic rat brain. Magn. Reson. Med. 49, 206–215 (2003).

36. Wu, D., Martin, L. J., Northington, F. J. & Zhang, J. Oscillating-gradient diffusion magnetic resonance imaging detects acute subcellular structural changes in the mouse forebrain after neonatal hypoxia-ischemia. J. Cereb. Blood Flow Metab. 39, 1336–1348 (2019).

37. Nilsson, M., Lasič, S., Drobnjak, I., Topgaard, D. & Westin, C.-F. Resolution limit of cylinder diameter estimation by diffusion MRI: The impact of gradient waveform and orientation dispersion. NMR Biomed. 30, (2017).

38. Paquette, M., Eichner, C., Knösche, T. R. & Anwander, A. Axon Diameter Measurements using Diffusion MRI are Infeasible. bioRxiv 2020.10.01.320507 (2021) doi:10.1101/2020.10.01.320507.

39. Palombo, M. et al. SANDI: A compartment-based model for non-invasive apparent soma and neurite imaging by diffusion MRI. Neuroimage 215, 116835 (2020).

40. Ianus, A., Alexander, D. C., Zhang, H. & Palombo, M. Mapping complex cell morphology in the grey matter with double diffusion encoding MR: A simulation study. Neuroimage 241, 118424 (2021).

41. Schiavi, S. et al. Dissecting brain grey and white matter microstructure: a novel clinical diffusion MRI protocol. bioRxiv 2022.04.08.487640 (2022) doi:10.1101/2022.04.08.487640.

42. Afzali, M. et al. Improving neural soma imaging using the power spectrum of the free gradient waveforms. in Proc. Intl. Soc. Mag. Reson. Med. 28 4426 (2020).

43. Jelescu, I. O., de Skowronski, A., Geffroy, F., Palombo, M. & Novikov, D. S. Neurite Exchange Imaging (NEXI): A minimal model of diffusion in gray matter with inter-compartment water exchange. Neuroimage 256, 119277 (2022).

44. Olesen, J. L., Østergaard, L., Shemesh, N. & Jespersen, S. N. Diffusion time dependence, power-law scaling, and exchange in gray matter. Neuroimage 251, 118976 (2022).

45. Olesen, J. L., Ianus, A., Shemesh, N. & Jespersen, S. N. Time dependence at ultra-high diffusion weighting reveals fast compartmental exchange in rat cortex in vivo. in Proc. Intl. Soc. Mag. Reson. Med. 30 1426 (2022).

46. Reynaud, O. Time-dependent diffusion MRI in cancer: tissue modeling and applications. Frontiers in Physics 5, 58 (2017).

47. Lampinen, B., Lätt, J., Wasselius, J., van Westen, D. & Nilsson, M. Time dependence in diffusion MRI predicts tissue outcome in ischemic stroke patients. Magn. Reson. Med. 86, 754–764 (2021).

48. Assaf, Y., Mayk, A. & Cohen, Y. Displacement imaging of spinal cord using q-space diffusion-weighted MRI. Magn. Reson. Med. 44, 713–722 (2000).

49. Jespersen, S. N., Olesen, J. L., Hansen, B. & Shemesh, N. Diffusion time dependence of microstructural parameters in fixed spinal cord. Neuroimage 182, 329–342 (2018).

50. Lee, H.-H., Papaioannou, A., Kim, S.-L., Novikov, D. S. & Fieremans, E. A time-dependent diffusion MRI signature of axon caliber variations and beading. Communications Biology 3, 1–13 (2020).

51. Aggarwal, M., Smith, M. D. & Calabresi, P. A. Diffusion-time dependence of diffusional kurtosis in the mouse brain. Magn. Reson. Med. 84, 1564–1578 (2020).

52. Xu, J. et al. Fast and simplified mapping of mean axon diameter using temporal diffusion spectroscopy. NMR Biomed. 29, 400–410 (2016).

53. Alexander, D. C., Dyrby, T. B., Nilsson, M. & Zhang, H. Imaging brain microstructure with diffusion MRI: practicality and applications. NMR Biomed. 32, e3841 (2019).

54. Novikov, D. S., Kiselev, V. G. & Jespersen, S. N. On modeling. Magn. Reson. Med. 79, 3172–3193 (2018).

55. Dell’Acqua, F. et al. Temporal Diffusion Ratio (TDR): A Diffusion MRI technique to map the fraction and spatial distribution of large axons in the living human brain. in ISMRM Conference Proceedings (2019).

56. Jelescu, I. O., Veraart, J., Fieremans, E. & Novikov, D. S. Degeneracy in model parameter estimation for multi-compartmental diffusion in neuronal tissue. NMR Biomed. 29, 33–47 (2016).

57. Callaghan, P. T., Jolley, K. W. & Lelievre, J. Diffusion of water in the endosperm tissue of wheat grains as studied by pulsed field gradient nuclear magnetic resonance. Biophys. J. 28, 133–141 (1979).

58. Panagiotaki, E. et al. Noninvasive quantification of solid tumor microstructure using VERDICT MRI. Cancer Res. 74, 1902–1912 (2014).

59. Roberts, T. A. et al. Noninvasive diffusion magnetic resonance imaging of brain tumour cell size for the early detection of therapeutic response. Sci. Rep. 10, 1–13 (2020).

60. Ianuş, A., Alexander, D. C. & Drobnjak, I. Microstructure imaging sequence simulation toolbox. in Simulation and Synthesis in Medical Imaging 34–44 (Springer International Publishing, 2016).

61. Callaghan, P. T. A simple matrix formalism for spin echo analysis of restricted diffusion under generalized gradient waveforms. J. Magn. Reson. 129, 74–84 (1997).

62. Drobnjak, I., Siow, B. & Alexander, D. C. Optimizing gradient waveforms for microstructure sensitivity in diffusion-weighted MR. J. Magn. Reson. 206, 41–51 (2010).

63. Drobnjak, I., Zhang, H., Hall, M. G. & Alexander, D. C. The matrix formalism for generalised gradients with time-varying orientation in diffusion NMR. J. Magn. Reson. 210, 151–157 (2011).

64. Duval, T. et al. Axons morphometry in the human spinal cord. Neuroimage 185, 119–128 (2019).

65. Aboitiz, F., Scheibel, A. B., Fisher, R. S. & Zaidel, E. Fiber composition of the human corpus callosum. Brain Res. 598, 143–153 (1992).

66. Rajkowska, G., Selemon, L. D. & Goldman-Rakic, P. S. Neuronal and glial somal size in the prefrontal cortex: A postmortem morphometric study of schizophrenia and Huntington disease. Arch. Gen. Psychiatry 55, 215–224 (1998).

67. Palombo, M., Alexander, D. C. & Zhang, H. Large-scale analysis of brain cell morphometry informs microstructure modelling of gray matter. in 29th ISMRM Annual Meeting and Exhibition (unknown, 2021).

68. Lackey, E. P., Heck, D. H. & Sillitoe, R. V. Recent advances in understanding the mechanisms of cerebellar granule cell development and function and their contribution to behavior. F1000Res. 7, (2018).

69. Houston, C. M. et al. Exploring the significance of morphological diversity for cerebellar granule cell excitability. Sci. Rep. 7, 1–16 (2017).

70. Tian, Q. et al. Comprehensive diffusion MRI dataset for in vivo human brain microstructure mapping using 300 mT/m gradients. Scientific Data 9, 1–11 (2022).

71. Ianuş, A. et al. Soma and Neurite Density MRI (SANDI) of the in-vivo mouse brain and comparison with the Allen Brain Atlas. Neuroimage 254, 119135 (2022).

72. Veraart, J. et al. Denoising of diffusion MRI using random matrix theory. Neuroimage 142, 394–406 (2016).

73. Dula, A. N., Gochberg, D. F., Valentine, H. L., Valentine, W. M. & Does, M. D. Multiexponential T2, magnetization transfer, and quantitative histology in white matter tracts of rat spinal cord. Magn. Reson. Med. 6 3, 902–909 (2010).

74. Grussu, F. et al. Neurite dispersion: a new marker of multiple sclerosis spinal cord pathology? Ann Clin Transl Neurol 4, 663–679 (2017).

75. Anaby, D. et al. Single- and double-Diffusion encoding MRI for studying ex vivo apparent axon diameter distribution in spinal cord white matter. NMR Biomed. 32, e4170 (2019).

76. Fick, R. H. J., Sepasian, N., Pizzolato, M., Ianus, A. & Deriche, R. Assessing the feasibility of estimating axon diameter using diffusion models and machine learning. in 2017 IEEE 14th International Symposium on Biomedical Imaging (ISBI 2017) 766–769 (2017).

77. Nunes, D., Cruz, T. L., Jespersen, S. N. & Shemesh, N. Mapping axonal density and average diameter using non-monotonic time-dependent gradient-echo MRI. J. Magn. Reson. 277, 117–130 (2017).

78. Ianus, A., Cruz, R., Chavarrias, C., Palombo, M., and Shemesh, N. Early microstructural aberrations in a mouse model of Alzheimer’s disease detected by Soma and Neurite Density Imaging. in Proc. Intl. Soc. Mag. Reson. Med. 30 2393 (2022).

79. Nilsson, M., Lätt, J., Ståhlberg, F., van Westen, D. & Hagslätt, H. The importance of axonal undulation in diffusion MR measurements: a Monte Carlo simulation study. NMR Biomed. 25, 795–805 (2012).

80. Lee, H.-H., Jespersen, S. N., Fieremans, E. & Novikov, D. S. The impact of realistic axonal shape on axon diameter estimation using diffusion MRI. Neuroimage 223, 117228 (2020).

81. Alves, R. et al. Correlation Tensor MRI deciphers underlying kurtosis sources in stroke. Neuroimage 247, 118833 (2022).

82. Budde, M. D. & Frank, J. A. Neurite beading is sufficient to decrease the apparent diffusion coefficient after ischemic stroke. Proc. Natl. Acad. Sci. U. S. A. 107, 14472–14477 (2010).

